# Large-scale Inference of Cell Lineage Trees and Genotype Calling from Noisy Single-Cell Data Using Efficient Local Search

**DOI:** 10.1101/2024.11.08.622704

**Authors:** Haotian Zhang, Yiming Zhang, Teng Gao, Yufeng Wu

## Abstract

In a multicellular organism, cell lineages share a common evolutionary history. Knowing this history can facilitate the study of development, aging, and cancer. Cell lineage trees represent the evolutionary history of cells sampled from an organism. Recent developments in single-cell sequencing have greatly facilitated the inference of cell lineage trees. However, single-cell data are sparse and noisy, and the size of single-cell data is increasing rapidly. Accurate inference of cell lineage tree from large single-cell data is computationally challenging. In this paper, we present ScisTree2, a fast and accurate cell lineage tree inference and genotype calling approach based on the infinite-sites model. ScisTree2 relies on an efficient local search approach to find optimal trees. ScisTree2 also calls single-cell genotypes based on the inferred cell lineage tree. Experiments on simulated and real biological data show that ScisTree2 achieves better overall accuracy while being significantly more efficient than existing methods. To the best of our knowledge, ScisTree2 is the first model-based cell lineage tree inference and genotype calling approach that is capable of handling datasets from tens of thousands of cells or more.

## Introduction

An adult human body contains about 30 trillion cells that perform various functions. These cells are interconnected via a shared cellular division history that spans development, aging, and disease. The evolutionary history of cells sampled from an organism can be depicted by a cell lineage tree, a fundamental model integral to the study of tumor evolution and developmental biology (Bizzotto et al., 2021; Coorens et al., 2021; Navin, 2014; Simeonov et al., 2021). A cell lineage tree is a rooted tree in which leaves are labeled by the extant cells and internal nodes represent progenitor cells that are ancestral to the extant cells. To simplify our notation, we use the terms “tree” and “cell lineage tree” interchangeably.

Reconstructing cell lineage trees from noisy single-cell DNA data has been actively studied in recent literature. Existing methods differ in their modeling assumptions and also in computational approaches. First, mutational models are essential for single-cell DNA data analysis, as mutations give rise to genomic variants, which constitute the primary data used for inferring cell lineage trees. The primary class of genomic variants studied in this paper is the single nucleotide variant (SNV). We do not consider more complex variants such as copy number variations (CNVs) for cell lineage tree inference (see the Discussion section). The infinite-sites (IS) model is a main mutational model for SNV variants. The IS model assumes that each mutant variant only arises *once* in the evolutionary history (i.e., there are no recurrent mutations). For example, in the cell lineage tree shown in Fig. 1(a), there is a single mutation for each SNV site. Many existing approaches assume the IS model (e.g., Jahn et al. (2016); Wu (2020); Kızılkale et al. (2022)). An alternative model is the finite-sites model (Zafar et al., 2017; Kozlov et al., 2022), which allows recurrent mutations at a genomic site.

**Figure 1.**
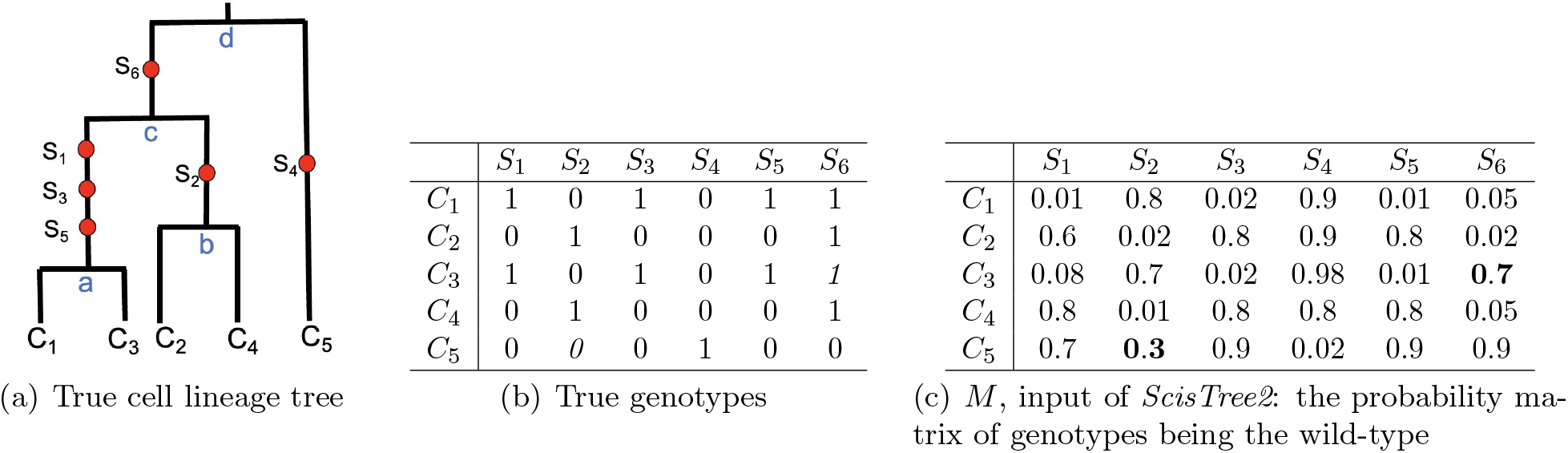
Illustration of cell lineage tree, genotypes with uncertainty and input of ScisTree2. Part 1(a): the true cell lineage tree with five cells and six sites (with mutations labeling branches). Infinite-sites model: one mutation per site. Four internal nodes: a to d. Part 1(b): the true (binary) genotypes. There are five cells and six SNV sites. Part 1(c): the input of ScisTree2is the genotype probability matrix of genotypes being the wild-type (0). The two boldfaced positions denote genotypes called using the maximum probability allele that do not agree with the true genotypes. For example, for the cell C_3_ and the site S_6_, M [3, 6] = 0.7, which would wrongly call this genotype the wild-type (0).

Existing cell lineage tree reconstruction methods also differ in high-level computational approaches. Several methods are based on probabilistic inference and use Markov chain Monte Carlo (MCMC) (Jahn et al., 2016; Zafar et al., 2017). A major downside of these methods is that they cannot handle large data. A more efficient alternative to MCMC is using *local search* to find optimal trees on a probabilistic model. Several newer methods including *ScisTree* (Wu, 2020) and *CellPhy* (Kozlov et al., 2022) adopt this approach. Lastly, there are parsimony-based approaches such as *HUNTRESS* (Kızılkale et al., 2022) which aims to make the fewest changes to genotypes so that the changed genotypes satisfy the IS model.

We previously developed a cell lineage tree inference method called *ScisTree* (Wu, 2020), which identifies the optimal tree that maximizes the posterior probability under the infinite-sites model through local search in tree space. *ScisTree* first constructs an initial tree from heuristically called genotypes using the well-known neighbor joining algorithm (Saitou and Nei, 1987). It then iteratively evaluates and identifies the tree with the highest posterior probability among those topologically similar to the current tree. Benchmarking on simulated data demonstrated that *ScisTree* accurately inferred trees and was significantly faster than several existing methods, including *SCITE* (Jahn et al., 2016).

However, *ScisTree* becomes inefficient when the number of cells exceeds 1,000. The size of single-cell data is rapidly increasing, both in the number of assayed cells and the amount of genomic information captured per cell. Single-cell genomic data with tens of thousands of cells is becoming available. For example, a recently developed variant caller, SComatic, can detect somatic SNVs from scRNA-seq and scATAC-seq, which typically contain 1,000s to 10,000s of cells (Muyas et al., 2023). Currently, no existing probabilistic cell lineage tree inference methods are capable of handling datasets of this size. Thus, a new method that can perform accurate lineage tree inference for large data is needed.

## Results

### *ScisTree2*: a software tool for efficient inference of cell lineage trees from large and noisy single-cell variation data

In this paper, we present *ScisTree2*, a new cell lineage tree inference approach that improves the original *ScisTree*. The input for running *ScisTree2* is generated by processing single-cell DNA-seq or other types of single-cell sequencing data with genetic variants. Given single-cell sequencing data (reads), the first step is using a genotype caller (e.g., GATK (McKenna et al., 2010)) to call the genetic variants from the single-cell sequence reads. Most genotype callers output a discrete probability distribution over possible genotypes for each cell and variant site. For simplicity, *ScisTree2* only considers two genotype states: wild-type (allele 0) and mutant (allele 1). *ScisTree2* takes the genotype probabilities of multiple single cells at multiple SNV sites and simultaneously infers the cell lineage tree and genotypes.

It is well known that single-cell data is noisy. One of the main sources of noise is allelic dropout (ADO), which is quite common in single-cell data and can lead to fewer or even no sequence reads at some SNV sites and cells. In current single-cell data, the ADO rate can be 50% or higher. ADO may lead to a wrongly called wild-type genotype while the true genotype is the mutant (i.e., producing a false negative error). Sequencing errors in single-cell DNA data are also common, which may lead to a wild-type allele being wrongly called a mutant (i.e., producing a false positive error). Noise in single-cell data implies that the called genotypes from single-cell data has significant *uncertainty*, which poses a major challenge for cell lineage tree inference.

*ScisTree* and *ScisTree2* take a genotype probability matrix *M* of size *n* rows by *m* columns as input. Here, *n* is the number of cells and *m* is the number of SNV sites. For each cell *c* and each site *s, M* [*c, s*] is equal to the *posterior* probability of the cell *c* having the *wild-type* genotype at the site *s* given the sequence reads at *s* (see Fig. 1(c)).

The original *ScisTree* (Wu, 2020) finds the tree *T* ^∗^ that gives the maximum posterior probability *P* (*T* ^∗^) *and* calls genotypes that fit the IS model using local search (Supplemental Methods, Sect. S1). Briefly, local search iteratively searches for an optimal cell lineage tree by finding the tree that gives the maximum posterior probability among all “neighboring” trees that are within a single tree rearrangement move, such as nearest neighbor interchange (NNI) or subtree prune and regraft (SPR), from the current tree. See the Methods section for more details. The original *ScisTree* performs the NNI search. The running time of the NNI local search is *O*(*Kn*^2^*m*) for NNI local search where *K* is the number of iterations which depends on how far the initial tree is from local optima (Supplemental Methods, Sect. S2). When data size is relatively large (say *n* = *m* = 1,000), *ScisTree* becomes slow.

The key technical contribution of the new *ScisTree2* approach is an SPR local search algorithm that runs in *O*(*Kn*^2^*m*) time, which is one order of magnitude faster than a naïve alternative. Moreover, the SPR local search is made more efficient by a branch and bound heuristic, which enables *ScisTree2* to infer cell lineage trees with 10,000 or more cells. See the Methods section for the details of the SPR local search algorithms. Experiments show that *ScisTree2* runs much faster than all existing cell lineage tree inference methods (except the classic distance-based methods such as neighbor joining). Moreover, *ScisTree2* outperforms existing methods in the accuracy of reconstructing the history of mutations and genotype calling.

### Simulated data

We compared *ScisTree2* with the following methods using simulated data: (i) *CellPhy* (Kozlov et al., 2022), (ii) *HUNTRESS* (Kızılkale et al., 2022), (iii) *SiFit* (Zafar et al., 2017), (iv) the original *ScisTree* (Wu, 2020) and (v) for small data only, *SCITE* (Jahn et al., 2016). In addition to comparing running time, we used the following metrics (all between 0 and 1, with 1 being the best) for accuracy comparison.

1. Genotype accuracy: the percentage of correctly called genotypes.
2. Tree accuracy: the percentage of the shared clades between the inferred trees and the true trees (which is equal to one minus the normalized Robinson-Foulds distance).
3. Ancestor-descendant (AD) *F* score (see, e.g., Kızılkale et al. (2022)): for the percentage of the pairs of mutations (*ma, mb*) where *ma* is ancestral to *mb* in the true tree but *not* so in the inferred tree.
4. Different-lineage (DL) *F* score (see, e.g., Kızılkale et al. (2022)): for the percentage of pairs of mutations (*ma, mb*) where *ma* is *not* ancestral to *mb* in the true tree but *ma* is ancestral to *mb* in the inferred tree.

Since *CellPhy* and *SiFit* are based on the finite sites model, we used the genotype caller implemented in *ScisTree2* to place the mutations on the trees inferred by *CellPhy* and *SiFit*. Then, we calculated the AD and DL accuracy based on the placed mutations. Note that this only affected AD and DL results for *CellPhy* and *SiFit*. Genotype accuracy of *CellPhy* and *SiFit* are based on the genotypes called by these two methods.

We used *CellCoal* (Posada, 2020) under the diploid infinite-sites model to simulate single-cell DNA sequence data. The simulation parameters (along with their default values) were set to: (i) the number of cells *n* (200), (ii) the number of SNV sites *m* (five times of *n*), (iii) the ADO rate (0.2), (iv) sequencing error rate (0.01), and (v) reads coverage (10x for high coverage data). For each settings of parameters, we generated 50 replicates and report the average of these replicates. We ran *CellPhy*-GL with the simulated VCF files from *CellCoal*. We ran *HUNTRESS* and *SCITE* with the maximum likelihood genotypes extracted from the VCF files. *ScisTree2* and *ScisTree* require posterior probability *Pr*(*G*|*D*) of genotype. Here, the data *D* are the sequence read counts for different alleles. *CellCoal* only calculates the likelihood *Pr*(*D*|*G*). We used Bayes’ Theorem to calculate *Pr*(*G*|*D*) from *Pr*(*D*|*G*). The details are given in the Supplemental Methods (Sect. S7).

### Simulated high-coverage data

#### Accuracy

Fig. 2 shows a comparison of the inference accuracy for different methods by varying the number of cells. Overall, *ScisTree2* and *CellPhy* had the highest tree accuracy, although *CellPhy* performed less well in genotype accuracy.

**Figure 2.**
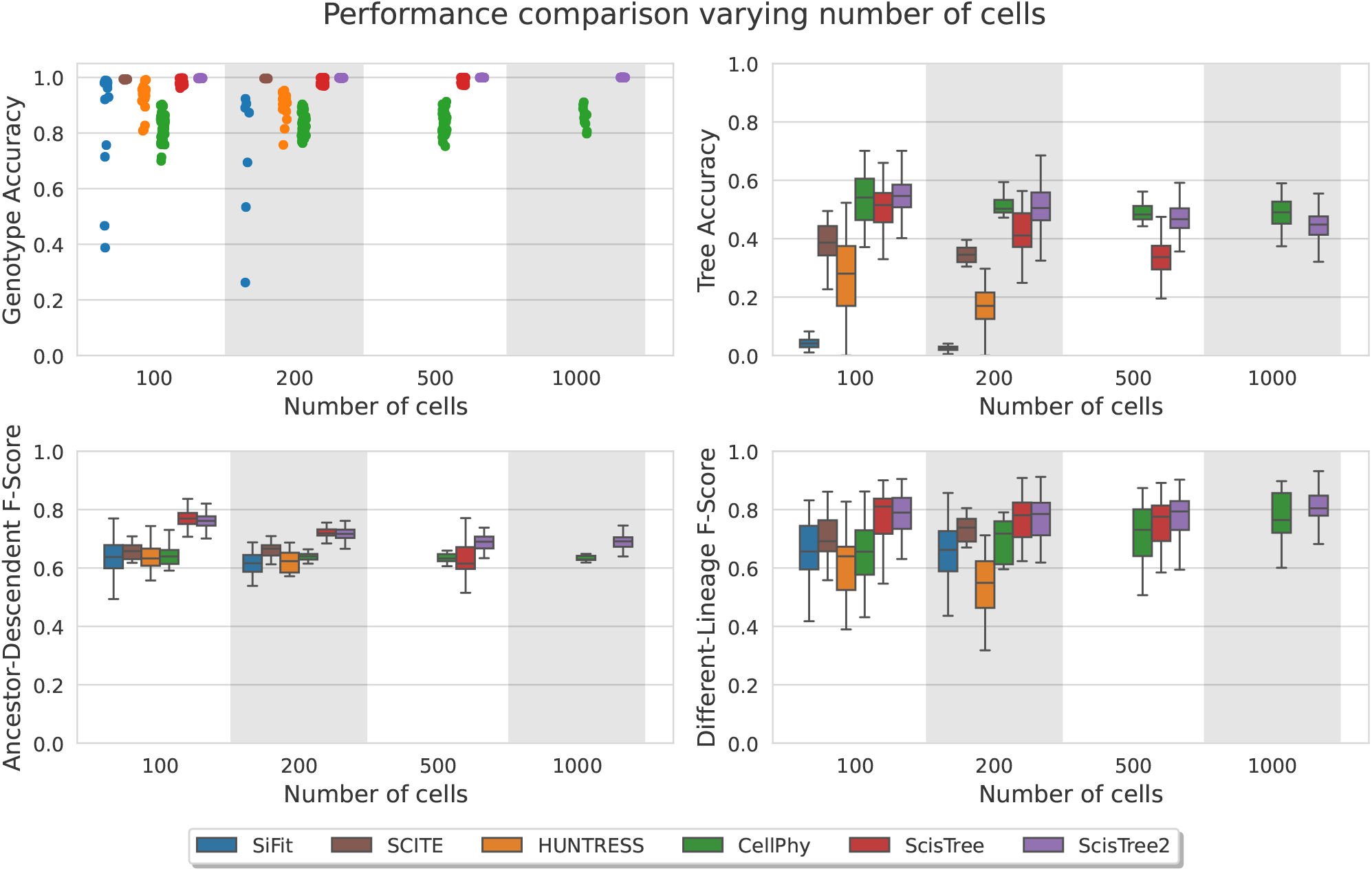
Comparison of inference accuracy of *ScisTree2* and five other methods: *SiFit, SCITE, HUNTRESS, CellPhy*, and the original *ScisTree* on simulated data with 10x coverage. The number of cells was varied in 100, 200, 500, and 1,000, the number of variants was set to 5× the number of cells, and methods that did not complete in one day are omitted. The Y-axis denotes four performance metrics: genotype accuracy, tree accuracy, AD score and DL score.

#### Running time

*SiFit, SCITE* and the original *ScisTree* are single-threaded while *HUNTRESS, CellPhy* and *ScisTree2* support multi-threading. The latter three were run using 30 threads. *CellPhy* was run on its “FAST” mode with the GT10 model that performs fast tree search from a single starting tree. *SiFit* was run with 10,000 iterations. *SCITE* was run with 500,000 MCMC iterations. The other methods used their default settings. All methods were tested on a Linux server with an Intel(R) Xeon(R) W-2195 CPU (36 cores). We tested four settings with varying numbers of cells and sites. The results are shown in Fig. 3. Results for methods that did not finish within a day are not reported. We report the CPU running time in Supplemental Table S1.

**Figure 3.**
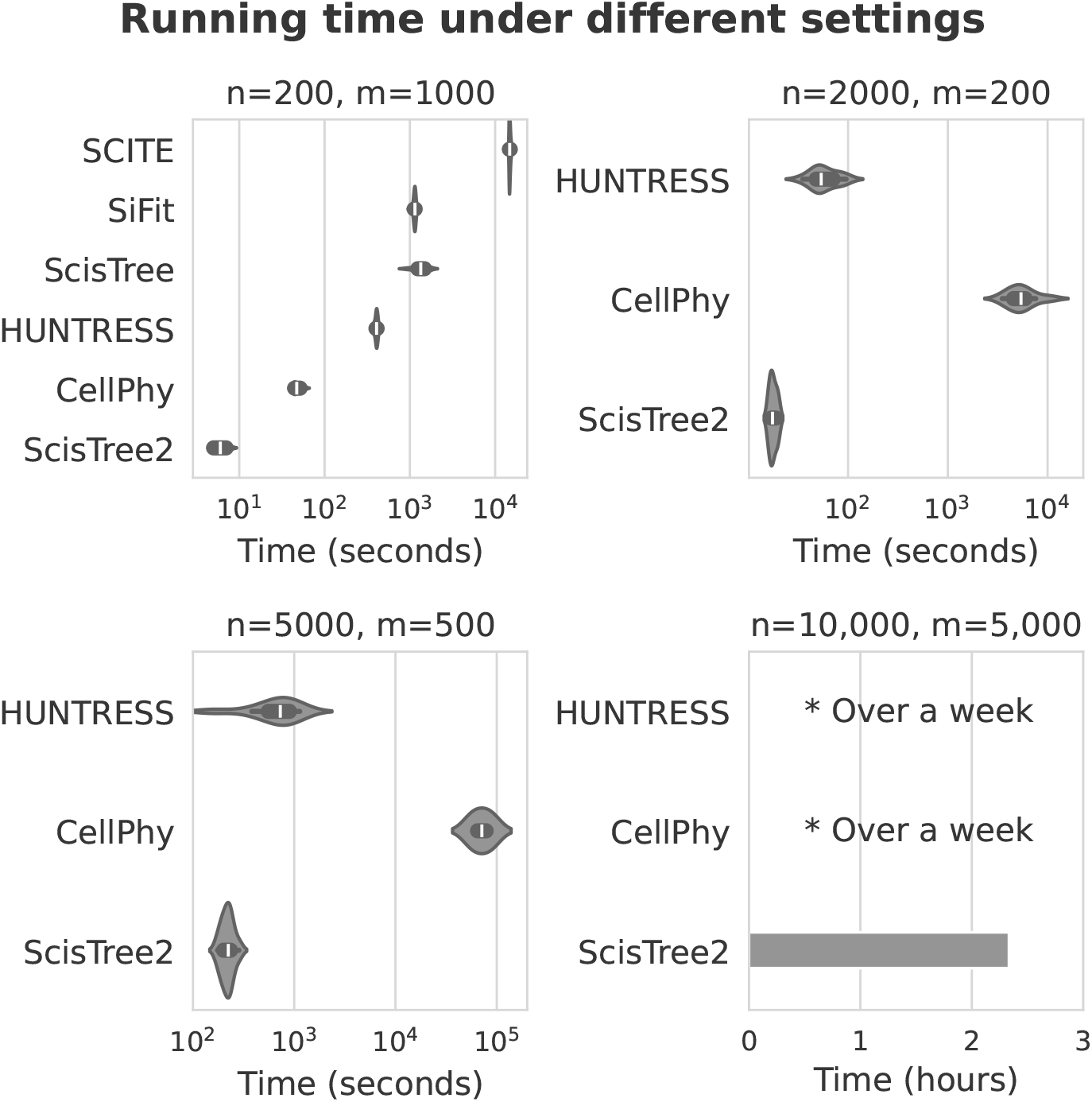
Comparison of the elapsed (user) running time between *ScisTree2* and other methods on simulated data with a varying number of cells (n) and sites (m), 30 threads were used for methods supporting multi-threading. Methods that are too slow are not reported.

Overall, *ScisTree2* demonstrated the highest computational efficiency, particularly for datasets with a large number of cells. *CellPhy* exhibited good scalability with respect to the number of sites but became computationally expensive as the number of cells increased. In contrast, *HUNTRESS* efficiently handled a large number of cells but experienced a substantial increase in runtime for datasets with many sites.

### Performance evaluation of *ScisTree2* and other methods by varying parameters on simulated data

We evaluated the effects of ADO rates and read coverage on each method using simulated data. Here, the number of cells was fixed to 200 and the number of sites is 1,000. The results are shown in Figs. 4 and 5. *ScisTree2* still outperformed other methods for data with high dropout rates or low coverage. These results demonstrated that *ScisTree2* performs well on data generated with various settings.

**Figure 4.**
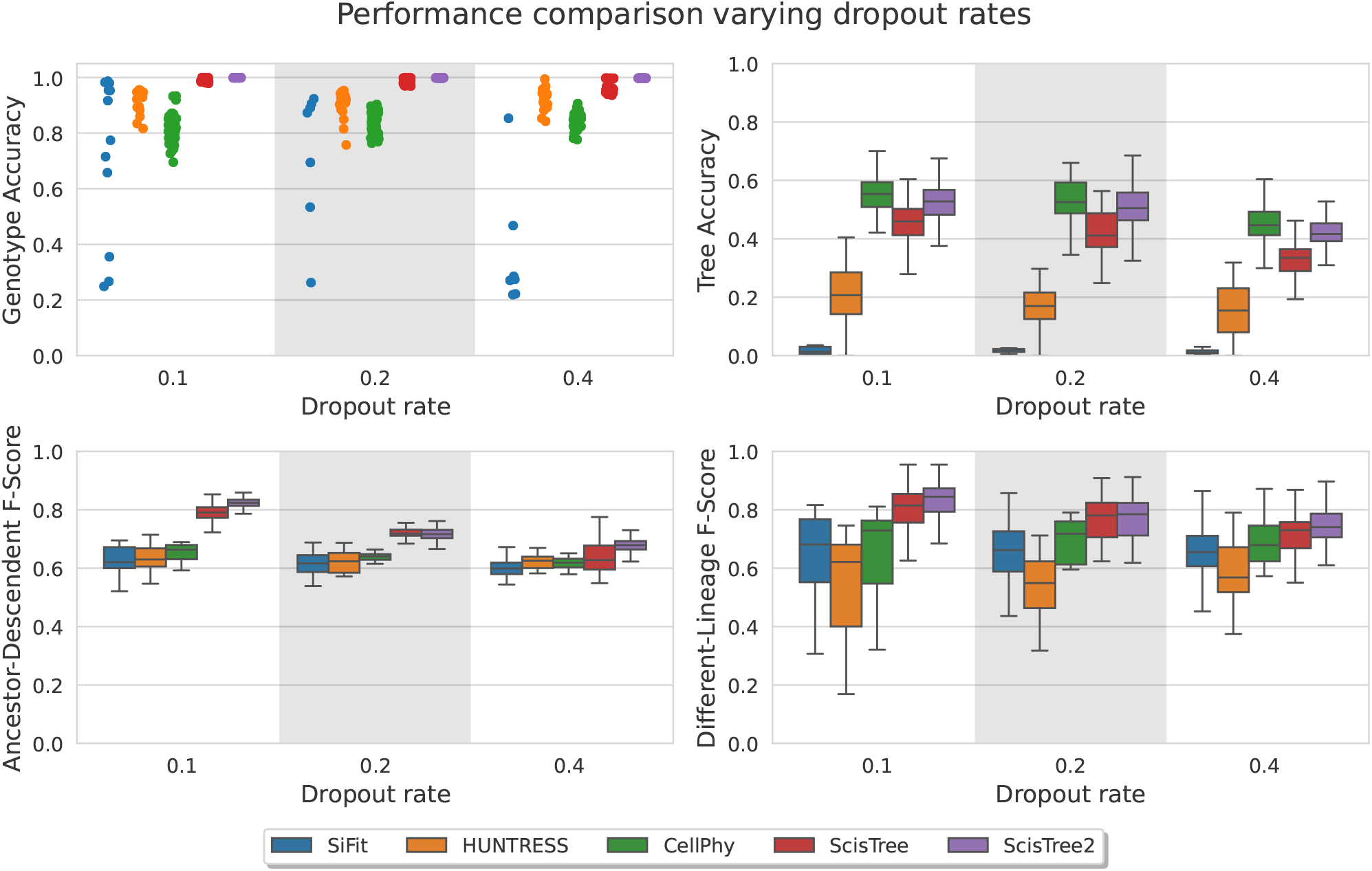
Performance comparison for *ScisTree2* and four other methods on various dropout rates. X-axis: dropout rates at 0.1, 0.2 and 0.4. Y-axis: performance metric.

**Figure 5.**
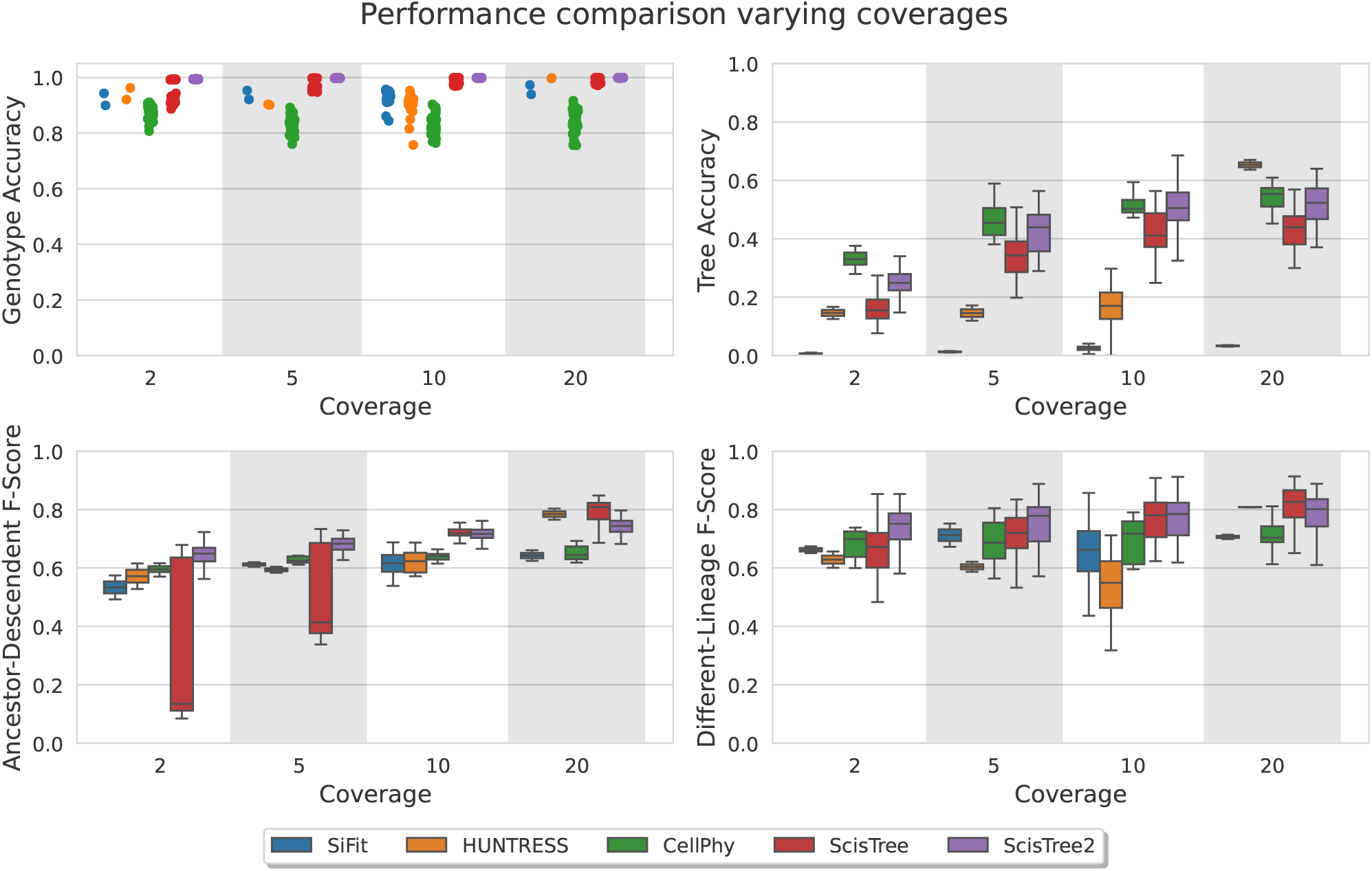
Performance comparison on various coverage (per site per cell). X-axis: coverage at 2x, 5x, 10x and 20x). Y-axis: performance metric.

### Accuracy of initial trees

We evaluated the accuracy of the initial trees constructed by the neighbor joining algorithm (Saitou and Nei, 1987). The results on the initial tree accuracy for fixed genotypes vs. uncertain genotypes are shown in Supplemental Fig. S1. Initial trees are reasonably accurate with high-quality data, but are less accurate than the trees inferred by *ScisTree2* for data with lower quality.

### Simulated low-coverage and low quality data

Some existing single-cell DNA sequence data have coverage lower than what we have simulated so far. For example, the 10X whole genome single-cell DNA sequence data can have as low as 0.01x coverage. Thus, it is useful to evaluate the performance of *ScisTree2* on data with coverage that is lower than 1x. Since *CellCoal* can only simulate data whose coverage is greater than 1x, to obtain data with very low coverage, we first simulated data with coverage 1x using *CellCoal*. Then we randomly dropped simulated reads with a certain probability. In this way, we simulated datasets with read coverages lower than 1x. The results are shown in Fig.6. Again, *ScisTree2* was highly accurate well on data with very low coverage.

**Figure 6.**
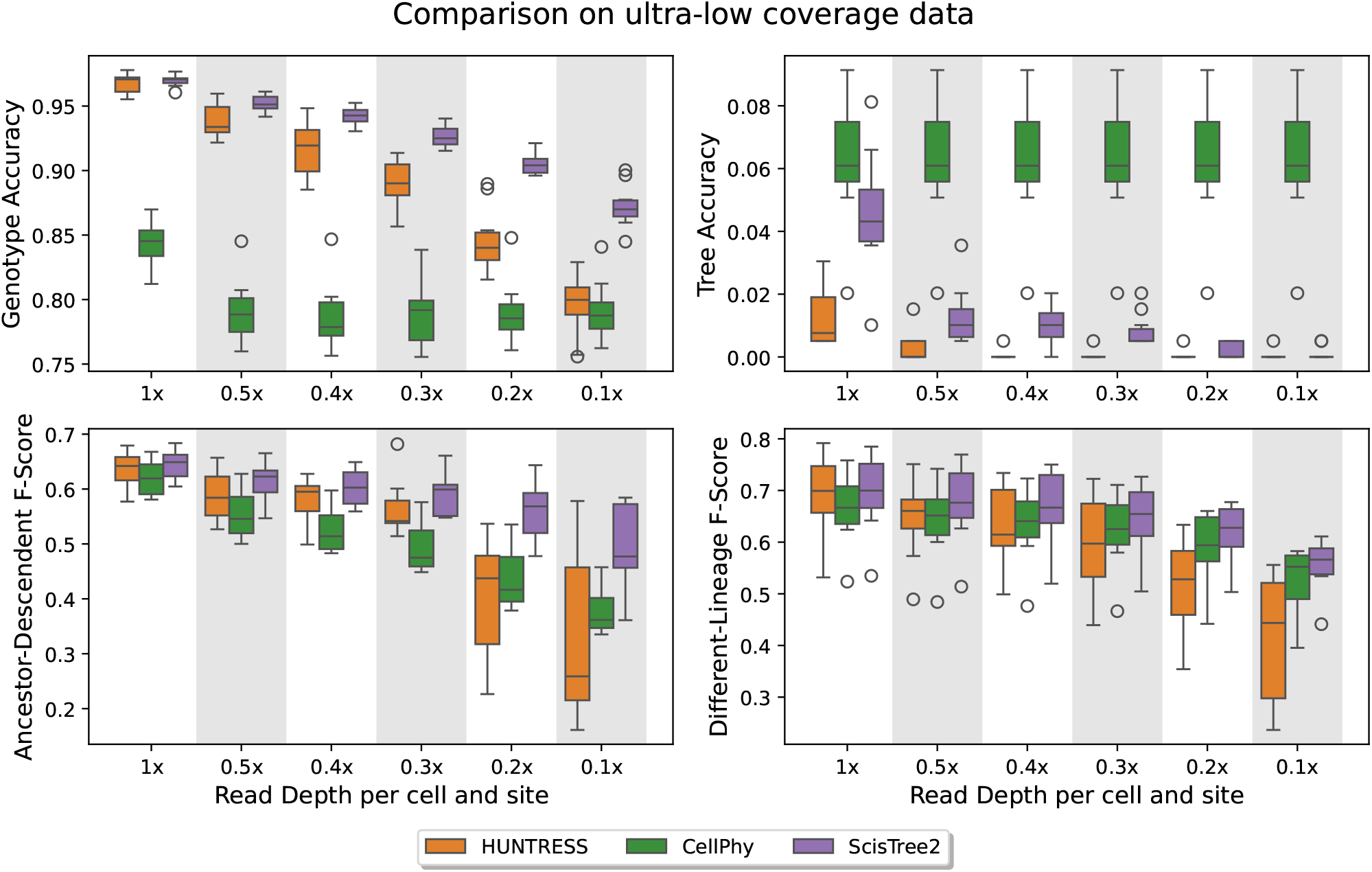
Performance comparison on data with lower than 1x coverage. 200 cells and 1,000 sites were used. *ScisTree2* outperformed other methods in genotype accuracy, AD and DL F-score: it called genotypes with more than 85% accuracy at 0.1x coverage. No methods achieved high tree accuracy.

In addition, we also simulated reads by manually adding noise (Supplemental Methods Sect. S10). The results in Supplemental Fig. S2 show that *ScisTree2* is robust to a certain level of noise.

### Simulation for targeted sequencing

Current single-cell targeted DNA sequencing usually generates data with a large number of cells but relatively small number of SNVs. To evaluate the performance of *ScisTree2* on data with similar settings as targeted single-cell sequencing, we simulated reads using *CellCoal* with 1,000, 2,000 and 5,000 cells while keeping the number of SNVs to be 500. *ScisTree2* clearly outperformed both *HUNTRESS* and *CellPhy* for genotype accuracy and AD/DL F scores for data with more cells than sites (Fig. 7). No method performed well in tree topology inference when the number of sites was small.

**Figure 7.**
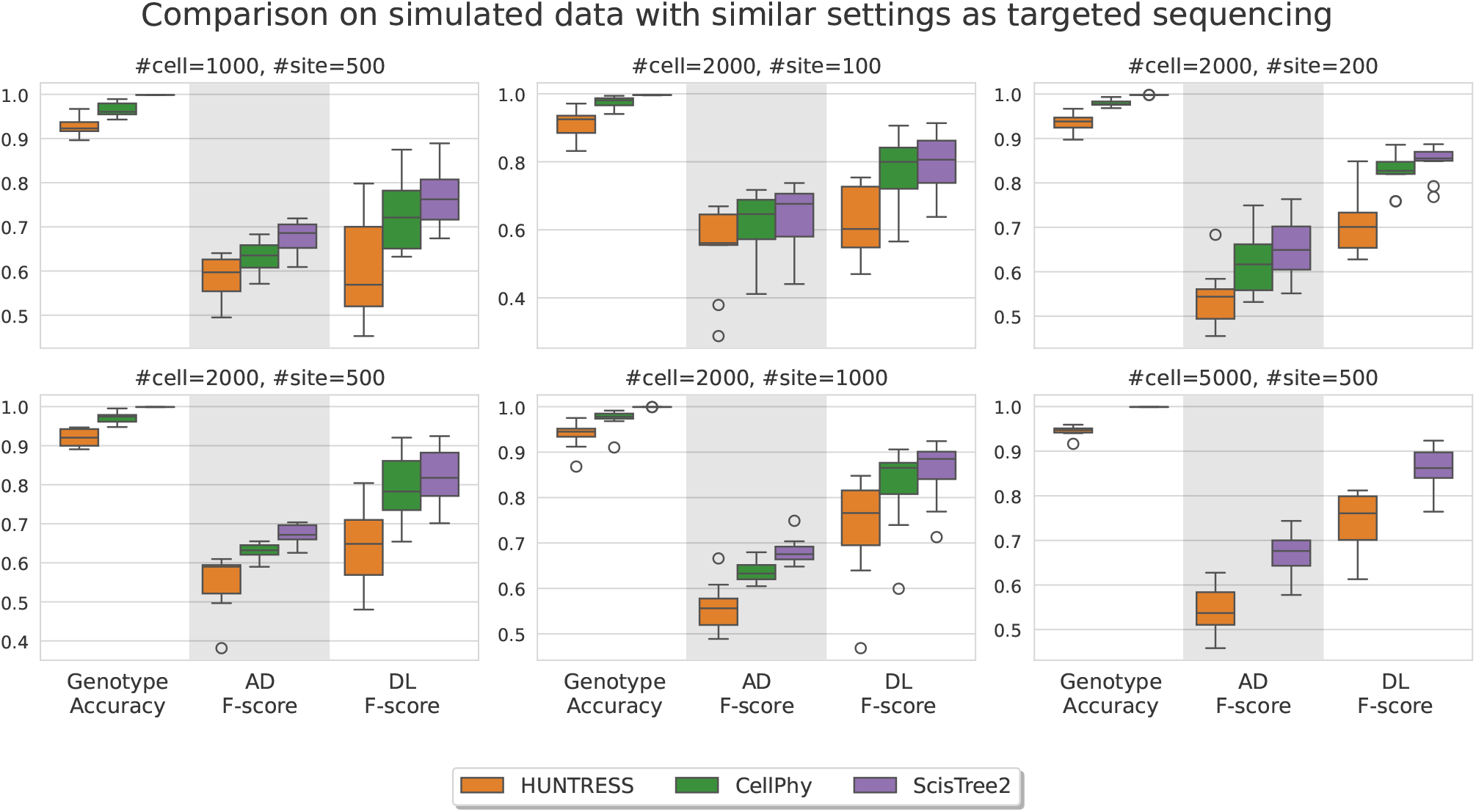
Comparison on simulated data with settings similar to targeted single-cell sequencing data with more cells and fewer sites. Tree accuracy is very low for all methods and is not shown.

### Simulation for data where the IS model does not strictly hold

*ScisTree2* along with *SCITE, HUNTRESS* and the original *ScisTree* assume the IS model. In real data, it is possible that the IS model does not strictly hold. That is, the data may contain both the sites that follow the IS model and the sites that follow the finite sites (FS) model. To evaluate the performance of methods on data that do not strictly fit the IS model, we performed simulations on data that are generated using different proportions of FS model sites (Supplemental Methods, Sect. S9). As shown in Fig. 8, the performance of all methods declined as the proportion of FS model sites increased. However, *ScisTree2* consistently outperformed other methods including *SiFit* and *CellPhy* that are specifically designed for the FS model. This suggests that *ScisTree2* is robust against minor violations of the IS model.

**Figure 8.**
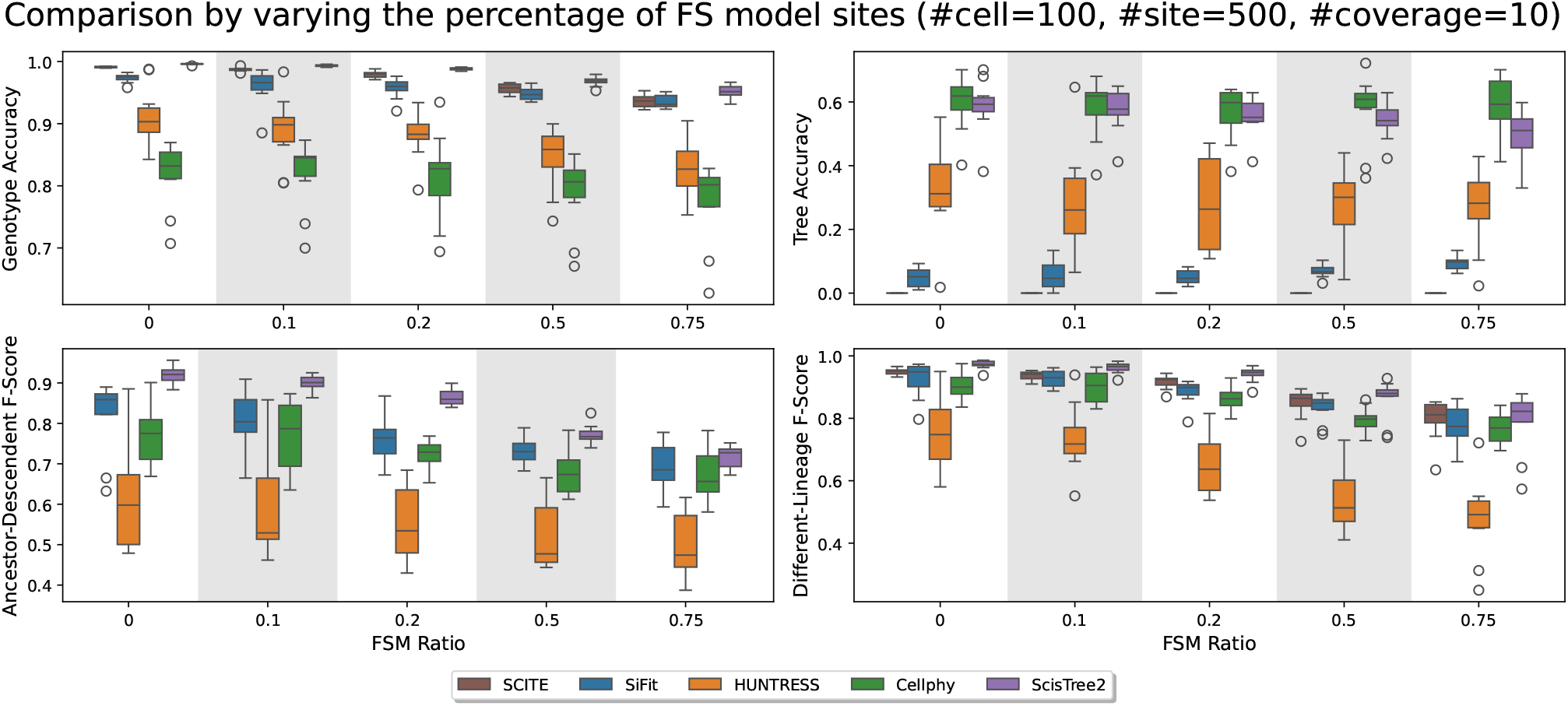
Comparison of simulated data with varying proportions of the FS model sites from 0 to 0.75. The dataset consists of 100 cells and 500 sites, with other parameters set to default. Note that AD and DL are calculated for sites that fit the IS model only.

### High-grade serous ovarian cancer single-cell DNA sequencing data

We evaluated the performance of *ScisTree2, CellPhy* and *HUNTRESS* on a low-coverage (with about 0.15x coverage) targeted sequencing dataset with 891 cells from three clonally related high-grade serous ovarian cancer (HGSOC) cell lines from the same patient that were processed by rigorous quality control (Laks et al., 2019). The clonal tree of nine clones for the HGSOC data in Laks et al. (2019) is shown in Fig. 9. The labels on the branches of the clonal tree are the genes where called mutations occur. Six genes, including three ancestral genes (*TP53, FOXP2* and *SUGCT*), and three clade genes (*HTR1D, INSL4* and *ZHX1*), are reported in Laks et al. (2019). Here, ancestral genes refer to the mutated genes shared by all tumor cells, and clade genes are those shared by a subset of clones. The true cell lineage tree for the HGSOC data is unknown, so we used the ordering of these six genes on the clonal tree (likely the most confident ones reported in Laks et al. (2019)) as the ground truth to test some aspects of inference accuracy. After dimensionality reduction and clustering, 891 cells were split into 9 clones based on copy number profiles, where the median clonal coverage was 15x (Laks et al., 2019). The resulting data contained the sequence read counts of 14,068 SNVs. We calculated the likelihood and posterior probabilities of the genotypes from the read counts (see the Supplemental Methods, Sect. S8). We only ran *ScisTree2* on the complete HGSOC data (with 14,068 SNVs), because *HUNTRESS* was slow on data with large number of sites and *CellPhy* was also slow which may be due to the large number of missing values in the data.

**Figure 9.**
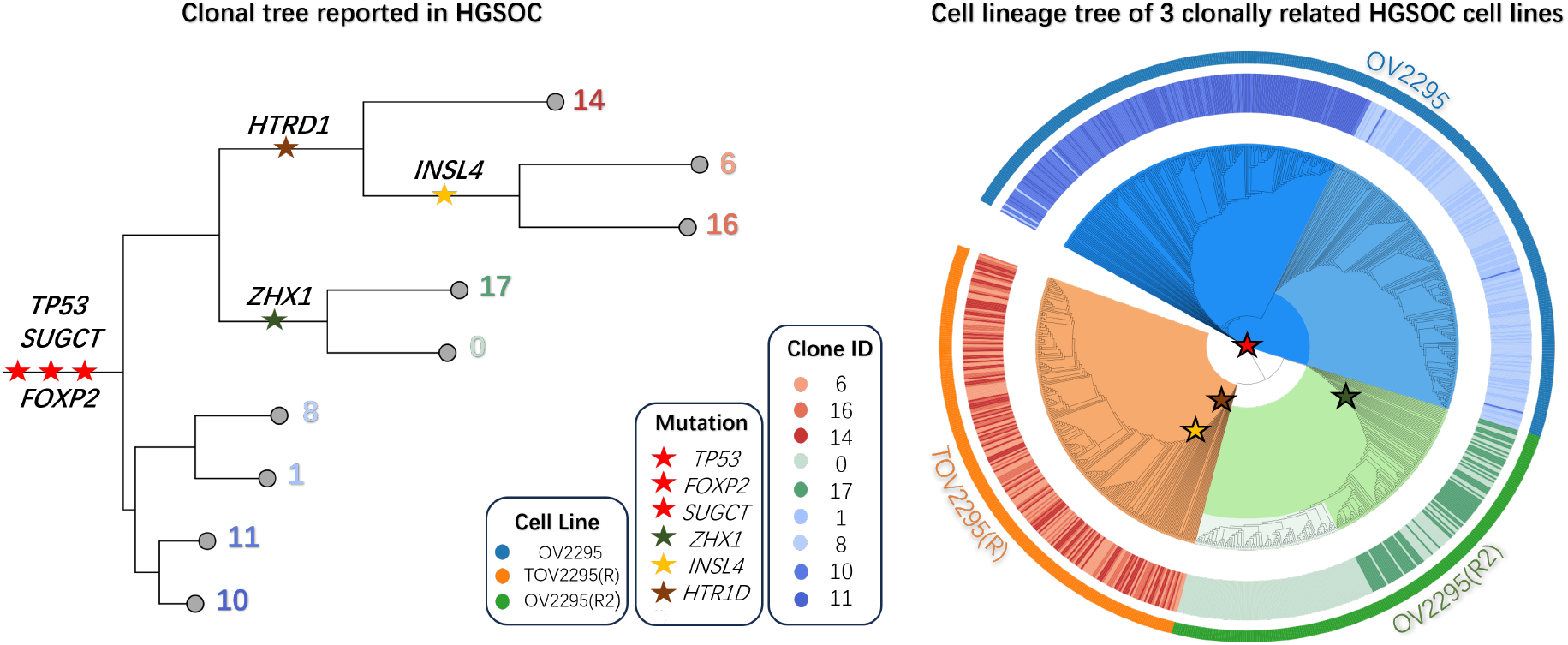
Analysis of the HGSOC data by *ScisTree2*. Left: the clonal tree of HGSOC with six mutated genes considered important for cancer development in the original study (Laks et al., 2019). Number at a leaf: clone ID (with a distinct color); Right: reconstructed cell lineage tree of tumor cells (colored w.r.t. their clones) with the same genes mapped on. Outer ring: 3 cell lines OV2295, TOV2295(R), and OV2295(R2). Inner ring: (colored) clones from the original study; Genes marked as red stars are ancestral genes, while others are clade genes.

#### Tree inference and mutated genes calling

We ran *ScisTree2* to infer the cell lineage tree and call genotypes for the complete HGSOC data. The called genotypes contained the 2,337 (16%) ancestral mutations that were shared by all 891 cells, 1,891 (14%) clonal mutations that were within a single clone, and 9,840 (70%) clade mutations that were located along branches of the clonal tree. Since genes affected by ancestral mutations may have a significant impact on early development of tumors, we used ANNOVAR (Wang et al., 2010) to annotate these ancestral mutations and find 48 exonic genes that may potentially be the key driver genes in ovarian cancer oncological pathways.

#### Evaluation of the called mutated genes and the inferred cell lineage tree

We used the inferred cell lineage tree to obtain the ordering of the six highlighted genes in Laks et al. (2019) and then compared our ordering with the gene ordering from the reported clonal tree (Fig. 9 left). The orderings of these six genes matched *perfectly* in the two studies. As expected, *TP53*, a commonly recognized cancer driver gene, appeared before all other mutated genes. In addition, we extracted 49 ancestral genes including the six reported genes based on the provided clonal tree in Laks et al. (2019) from the original study. All 48 genes found by *ScisTree2* were among them, with only one gene, *XXYLT1*, which was not identified as an ancestral gene in *ScisTree2*. We also investigated the topological concordance between the inferred cell lineage tree (Fig. 9 right) and the clonal tree in Laks et al. (2019). Each of the three cell lines formed a distinct clade in the cell lineage tree as expected. Within each cell line, the inferred cell lineage tree agreed with the clonal tree for the cell line OV2295 and OV2295(R2), but differed in many parts for cell line TOV2295(R). The clonal tree in Laks et al. (2019) was constructed by first clustering the cells into clones using copy number variations and then building trees using SNVs and breakpoints. Our results show that *ScisTree2* may be useful in clonal analysis with only SNV variants.

#### Analysis of a reduced HGSOC data

To compare with *HUNTRESS* and *CellPhy*, we constructed a smaller subset from the HGSOC data. We followed the same filtering approach used in Kızılkale et al. (2022). We filtered out sites with more than 650 missing values, which led to a reduced-size dataset with only 789 SNV sites. The original study in Laks et al. (2019) mapped these SNVs to the branches of the clonal tree, which we used as the ground truth for comparing different methods. *ScisTree2* needs two hyperparameters: the ADO rate and genotype priors, which were not known for the HGSOC data. To test the effects of these parameters, we show results with multiple settings of these parameters (Supplemental Methods Sect. S10). We collected the AD/DL F scores and the running time of *ScisTree2*, the neighbor joining algorithm (implemented in *ScisTree2*), *HUNTRESS* and *CellPhy*. The results are shown in Fig. 10. In general, *ScisTree2* outperforms the other methods for most settings in these data. Moreover, our results indicate that *ScisTree2* is robust to the ADO rate settings, and an appropriately chosen prior improves overall inference accuracy (Supplemental Fig. S3). Therefore, we recommend using genotype priors computed from allele frequencies as the default setting.

**Figure 10.**
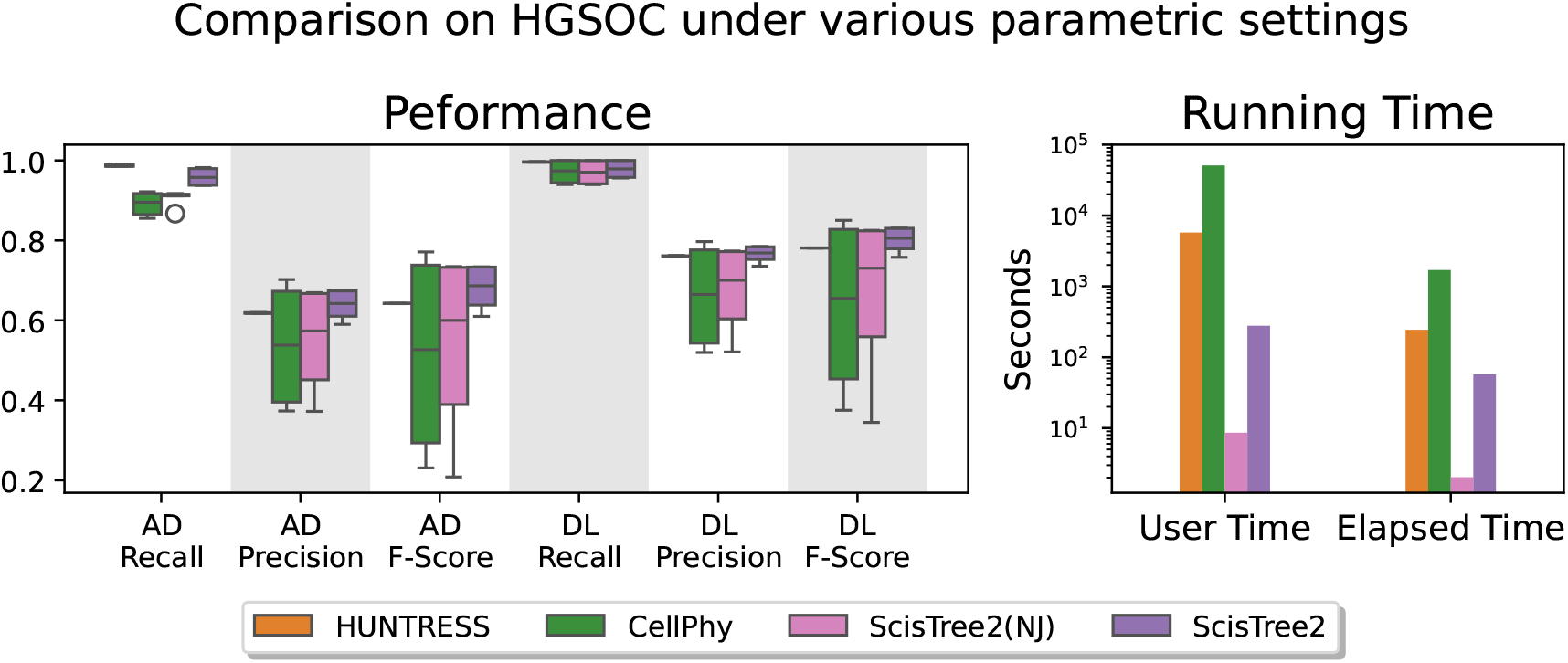
A comparison of *ScisTree2* with *HUNTRESS, CellPhy* and neighbor joining on a reduced HGSOC data. Left: AD and DL F scores of four methods for mutation ordering. Results from runs with different settings of parameters (e.g., ADO rates) are reported. Right: running time (in seconds) for user (CPU) time and elapsed (wall clock) time.

## Discussion

### Integrated analysis of single cells from multiple clones vs. analysis of single clones

A commonly used approach for analyzing large single-cell data is first clustering the cells into clones and then analyzing the cells from a single clone (Leung et al., 2017). This approach takes advantage of existing single-cell clustering methods and avoids the computational burden of analyzing large single-cell data. However, a main disadvantage of this approach is that errors can potentially be introduced in the clustering step. Because single-cell data usually contains significant noise, clustering single cells is not trivial. The efficiency of *ScisTree2* may allow integrated analysis of cells from multiple clones and can potentially avoid clustering errors. As shown in the Results Section, the inferred cell lineage tree can provide the clustering of cells from different cell lines. More experiments will be needed to compare the performance of integrated analysis and single-clone analysis.

### The infinite-sites model

One reason for the efficiency of *ScisTree2* is its assumption of the infinite-sites (IS) model. The IS model greatly simplifies the underlying probabilistic model of *ScisTree2*. While the IS model is popular in the cell lineage tree inference literature, the IS model may not hold for some data. As we show in the Results section (Fig. 8), *ScisTree2* is still reasonably accurate when the deviation from the IS model in the data is moderate. Nonetheless, it is useful to consider extending the model of *ScisTree2* to allow violations of the IS model. Since the IS model appears to perform reasonably well in practice, it may be prudent to consider using models that allow moderate deviations from the IS model. In the literature, Kuipers et al. (2022) explored extensions of the IS model to allow some recurrent mutations as well as mutation losses. We note that such extensions can lead to more complex probabilistic models and less efficient inference approaches. Thus, there may be a trade-off between accuracy and efficiency for inference methods.

### Beyond binary single nucleotide variants

In this paper, we assume that SNV genotypes are binary. In contrast, ternary SNV genotypes are allowed in Jahn et al. (2016); Wu (2020). We note that if the IS model holds strictly, SNV genotypes should be binary when recombination is absent. Ternary genotypes are only possible with additional mutational events, e.g., recurrent mutations or deletions of wild-type alleles at SNV sites. More generally, when copy number aberrations occur, single-cell genotypes of SNVs may no longer be limited to be binary or ternary. We note that the efficiency of *ScisTree2* is based on the simplicity of binary SNV data and the underlying mutation model. Some generalization is possible. For example, Wu (2020) shows that probabilities of ternary genotypes can be calculated efficiently with additional assumptions on the mutation model. Extending the *ScisTree2* approach to support more general single-cell genotypes and mutation models remains a research question.

### Applying *ScisTree2* in cancer genomics analyses

Inference of the cell lineage tree is very relevant to single-cell cancer genomics. For example, finding clonal populations is a common problem in cancer genomics. A clonal population corresponds to a clade (subtree) in the cell lineage tree. While *ScisTree2* does not find clonal populations at present, *ScisTree2* calls the genotypes and also places mutations on the tree. Such information may be useful for developing applications for downstream analyses in cancer genomics.

The ability of *ScisTree2* to analyze a large number of cells may be useful in large-scale cancer genomics analyses. One such analysis is identifying *rare* cancer subclones, which can drive disease recurrence and therapy resistance. For example, relapse of acute myeloid leukemia can be seeded by a *minor subclone* that represents approximately 1% of the primary tumor (Wong et al., 2015). Such clinically-relevant subpopulation can only be identified by analyzing thousands of cells in the tumor.

### Speeding up SPR local search

Since local search is a common strategy for phylogeny inference, speeding up SPR local search has been previously studied in the literature. For example, Hordijk and Gascuel (2005) applied various heuristics to speed up the local SPR search for general phylogeny inference. While such heuristics can achieve significant speedup in practice, there is little theoretical guarantee about the achieved speedup. In contrast, SPR local search algorithm in *ScisTree2* has proven a theoretical speedup of one order of magnitude. Such speedup is made possible by the *ScisTree2*’s probabilistic model. This simplified model leads to a much simpler structure in its probability function than the one used in the general maximum likelihood tree inference. Working with simpler but still reasonably accurate probabilistic models may be the key for speeding up cell lineage tree inference for large data.

## Methods

### The original *ScisTree* method

To present the new *ScisTree2* method, we start by briefly explaining the key aspects of the original *ScisTree* method. See the Supplemental Methods (Sections S1 and S2) for a more detailed discussion of the *ScisTree* method.

Compared to *CellPhy, ScisTree* uses a simpler probabilistic model, which is designed for the IS model. *ScisTree* aims at maximizing the **posterior** probability *Pr*(*G*|*D*) for the given sequence reads *D* and some genotypes *G* among all genotypes G that satisfy the IS model. We say that *G* satisfies the IS model if there exists a potentially multifurcating phylogeny *T* (called a *perfect phylogeny* (Gusfield, 1991)) where leaves are labeled by the cells (rows) in *G*, and each column (site) *c* of *G* labels a single branch of *T* such that exactly the cells below this branch are the mutants of *c*. See Figs. 1(a) and 1(b) for an illustration.

The posterior probability *Pr*(*G*|*D*) is calculated as follows. We let *G*(*c, s*) be the (binary) genotype of the individual cell *c* at the site *s*. Note that we also use the term genotype to refer to the list of genotypes at multiple sites or for multiple cells. Then,

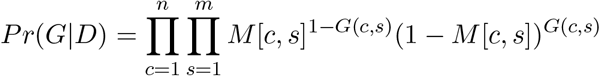

where *G* satisfies the IS model, *n* is the number of cells and *m* is the number of SNV sites. Here, we assume that each genotype at a site *s* for a cell *c* is independent. Recall that *M* is the genotype probability matrix of size *n* by *m* that is obtained from *D* during genotype calling. See the Supplemental Methods (Sect. S1) for a more detailed discussion of this probabilistic model.

Finding *G*^∗^ = arg max*G Pr*(*G*|*D*) for the given genotype probability matrix *M* is known to be NP complete (Wu, 2020). A key insight made in *ScisTree* is that if the underlying rooted and binary tree *T* is *given*, then *G*^∗^ can be found in *O*(*nm*) time by finding the best branches to place mutations on the given *T* as follows. Recall that the set of mutants of a site *s* must form a *single* clade (subtree) in *T*. Then, for each branch ending at node *v* in *T*, we define *Qs*(*v*) as

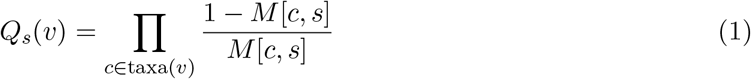

where taxa(*v*) is the set of leaves below *v*. Recall that *M* [*c, s*] denotes the probability of the wildtype genotype of the cell *c* and the site *s. Qs*(*v*) only concerns the cells within the subtree rooted at *v*, and has a simple product form. Let *v*1 and *v*2 be the two children of *v*. Then, *Qs*(*v*) = *Qs*(*v*1)*Qs*(*v*2). Thus, we can calculate all *Qs*(*v*) in *O*(*n*) time for each site in a bottom-up way using dynamic programming. We denote the set of nodes of *T* as nodes(*T*). To obtain the maximum posterior probability at a site *s*, we place the mutation for *s* on the branch whose destination node is *v*^∗^ where *v*^∗^ = max_*v*∈nodes(*T*)_ *Qs*(*v*). The maximum posterior probability *Pr*(*G*^∗^|*T, D*) of genotypes conditional on *T* is then:

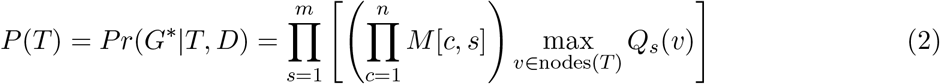

**Example** Note that when placing mutations on a given tree, each site is treated independently of other sites. So we use the site *S*1 in Fig. 1(c) to show how to find where its mutation is located on the tree in Fig. 1(a). We write *Q*1(*v*) as *Q*(*v*) here to simplify the notation. At leaves, 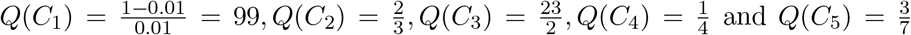. For internal nodes, 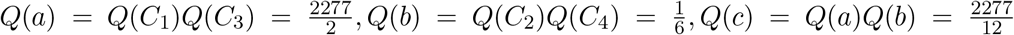, and 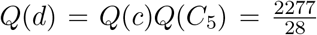. We place the mutation for *S*1 at the branch right above the node *a* because *Q*(*a*) is the *largest* among all the *Q* values for *S*1.

### Local Search

Local search is a popular phylogenetic inference approach. At a high level, local search implemented in *ScisTree* and *ScisTree2* works as follows.

1. Construct an initial rooted binary tree *T*0 based on ℳ. Initialize *T*opt ← *T*0, *P*opt ← *P* (*T*0) and *G*opt ← *G*0.
2. Find rooted binary trees 𝒯*c* that are within one tree arrangement operation (NNI or SPR) from *T*opt.
3. Let *T* ∈ 𝒯 *c* that maximizes the posterior probability *P* (*T*) for some genotypes *G*. If *P* (*T*) ≥ *r*opt∗*P*opt (*r*opt is a fixed constant that is slightly larger than 1.0), set *T*opt ← *T, P*opt ← *P* (*T*), *G*opt ← *G* and go to step 2. Otherwise, stop.

See the Supplemental Methods (Sect. S2) for more details on the original *ScisTree* method.

### Efficient SPR local search

We call the set of trees that can be obtained by a single tree rearrangement operation (e.g., NNI or SPR) as the 1-operation neighborhood of the current tree. The original *ScisTree* only implements the NNI local search. The size of 1-NNI neighborhood of a tree *T* with *n* taxa is *O*(*n*). A main disadvantage of the NNI local search is that the 1-NNI neighborhood is rather restricted. This makes NNI local search easily get trapped in local optima. An alternative to NNI is (rooted) subtree prune and regraft (SPR), which is more commonly used in phylogenetic inference. An SPR operation prunes (i.e., detaches) a subtree of a tree *T* and regrafts (i.e., re-attaches) to another branch of *T*. The size of 1-SPR neighborhood is *O*(*n*^2^), and contains the 1-NNI neighborhood as a subset. Naïvely, one can calculate the likelihood for each of the *O*(*n*^2^) trees in the 1-SPR neighborhood. But this leads to *O*(*n*^3^*m*) running time for *n* cells and *m* SNV sites, because the time for finding the maximum posterior probability for a single tree is *O*(*nm*). This is too slow even for data of moderate size. We now present a novel algorithm that finds the best tree in the 1-SPR neighborhood in *O*(*n*^2^*m*) time.

#### High-level idea

We focus on computing the posterior probability *P* (*T* ^′^) where *T* ^′^ is obtained by a *single* SPR operation on the current tree *T*. This is illustrated in Fig. 11. By Equation 2, we need to find the maximum over all the *Q* values in *T* ^′^. The key observation is that we do not need to calculate all *Q* values of *T* ^′^ from scratch. This is because *T* and *T* ^′^ are very similar topologically. Almost *all* the nodes in *T* are also in *T* ^′^. At these shared nodes, *Q* values in *T* ^′^ either are the same as those in *T*, or can be easily computed from the *Q* values in *T*. Careful consideration of the relations between the *Q* values in *T* and *T* ^′^ leads to an efficient algorithm to find the largest value *Q* in *T* ^′^, after some pre-processing is performed on *T*. We now give the details of this algorithm.

**Figure 11.**
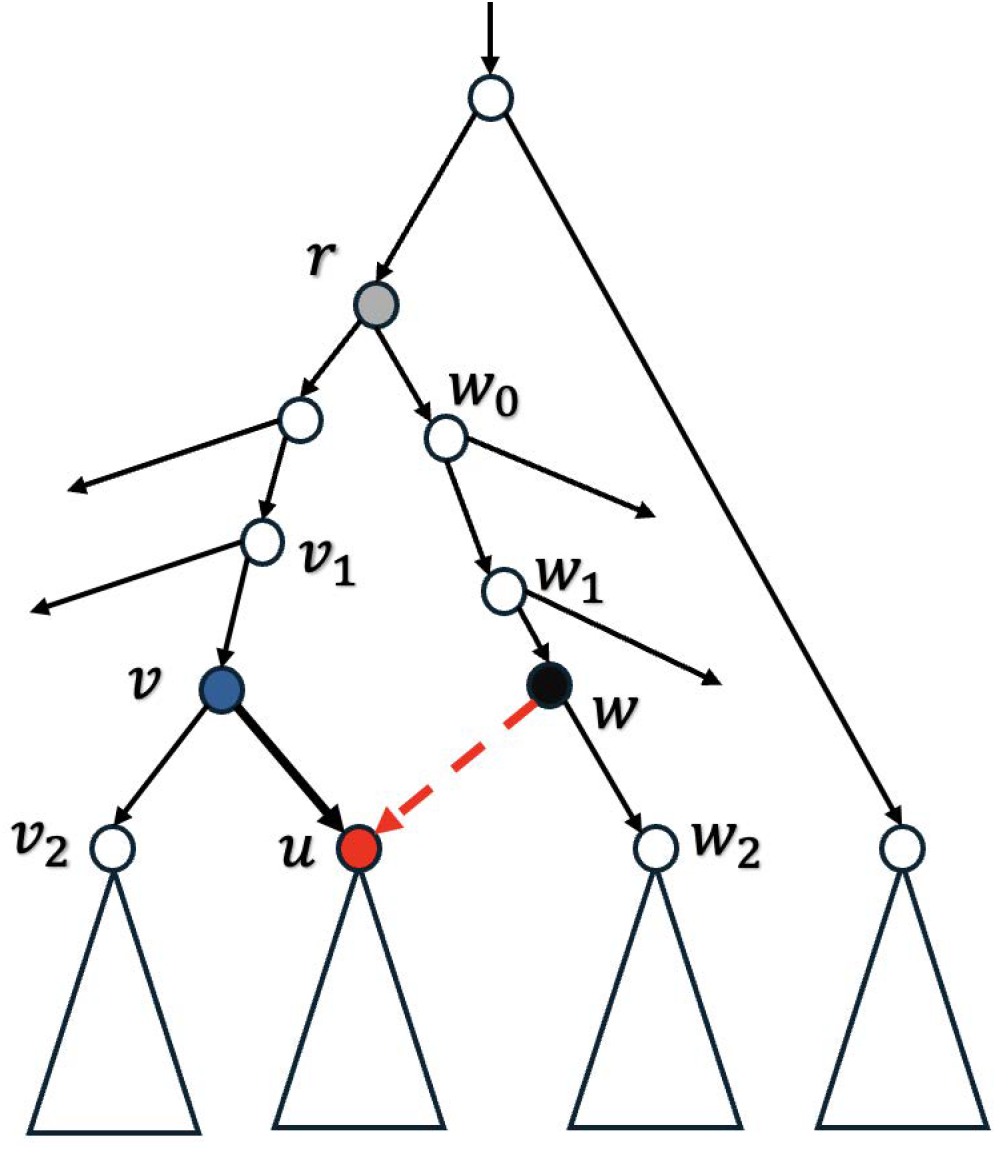
An illustration of (rooted) SPR local search. The subtree rooted at *u* (whose parent is *v*) is pruned and regrafted to the edge entering *w*2. Note: the lowest common ancestor of *v* and *w*2 in the tree before SPR is *r*.

We consider the SPR operation in Fig. 11 that prunes a subtree *Tu* (rooted at the node *u*) of *T* and regrafts so that the parent of *u* is a new node *w* in *T* ^′^, which breaks the edge (*w*1, *w*2) into two edges. We let *v*2 be the sibling of *u* and *v*1 be the parent of *v* in *T*. We let node *r* be the lowest common ancestor (LCA) of *v* and *w*2 in *T*. Recall that the posterior probability is the product of the maximum *Qs*(*a*) (over each node *a* of *T*) times a constant factor for each site *s* (Equation 2). To simplify the notation, we omit the subscript *s* by writing *Qs*(*a*) as *Q*(*a*) throughout this paper with the understanding that *Q*(*a*) is for a specific site. We denote *Q*^′^(*x*) to be the updated *Q*(*x*) value for the node *x* on the new tree *T* ^′^ after the SPR move shown in Figure 11.

We now show that we can find the maximum of *Q*^′^(*x*) values efficiently *without* calculating all *Q*^′^(*x*) values explicitly. Note that *T* ^′^ has a new node *w* that is not in *T*, while *v* is in *T* but not in *T* ^′^. All the other nodes are the same in *T* and *T* ^′^. Recall that *Q*(*a*) is equal to the product of the ratios between the probability of the taxon *t*(*b*) being the mutant (allele 1) and that of being the wild-type (allele 0) for each leaf *b* within the subtree *Ta* (Equation 1). A path from the node *u* to the node *v* of a tree *T* is denoted as *u* → *v*, which consists of all nodes from *u* to *v* sequentially, and *u* and *v* are in the path unless otherwise stated. We have the following observation. For any node *x* ∈ *T* ^′^, exactly one of the following four cases holds for *Q*^′^(*x*):

1. *x* is a node in *T* that either (i) is *not* on the paths *r* → *v* and *r* → *w*1, or (ii) *x* = *r*. Then, *Q*^′^(*x*) = *Q*(*x*). This is because the SPR move does not change the set of taxa under node *x* and thus *Q*^′^(*x*) is equal to the product of the same set of ratios as in *Q*(*x*).
2. *x* ≠ *r* and is on the path 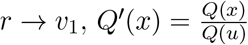. This is again because *Q*(*x*) is the product of ratios of probabilities of genotypes within *Tx* being 1 vs. 0; after the SPR move, the leaves under *u* are *removed* from 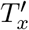 and thus are *absent* from the product for *Q*^′^(*x*).
3. *x* ≠ *r* and is along the path *r* → *w*1, *Q*^′^(*x*) = *Q*(*x*)*Q*(*u*). This is because now the leaves under *u* are *inserted* in 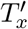.
4. *x* = *w* (i.e., *x* is the only new node in *T* ^′^ added by SPR). Then *Q*^′^(*w*) = *Q*(*w*2)*Q*(*u*).

We now perform some preprocessing steps on *T* that allow the evaluation of *multiple* nodes together that fall into the same case as listed above. The preprocessing is performed only once for the current tree *T* before the iteration of local search starts from *T*.

### Preprocessing of the original *T*

We have the following definitions for *T*. For each node *a* of *T*, 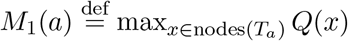. That is, *M*1(*a*) is the maximum value of *Q*(*x*) over all nodes *x* within the subtree *Ta* (it is allowed *x* = *a*). For a leaf *a*0, *M*1(*a*0) = *Q*(*a*0). For an internal node *a* with two children *a*1 and *a*2,

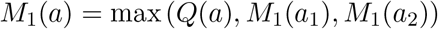

Note that the *M*1(*a*) values are for the nodes *a* in the original *T before* the SPR move. Recall that the *Q*(*a*) values are already computed. Therefore, all *M*1(*a*) values can be computed in *O*(*n*) time by performing a bottom-up traversal of *T*.

We denote the parent of node *b* in *T* as *p*(*b*). For each node *a* and a node *b* ≠ *a* where *b* ∈ nodes(*Ta*) (i.e., *b* is within the subtree *Ta*), define 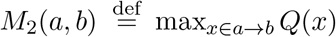. That is, *M*2(*a, b*) is the maximum value of *Q*(*x*) over all nodes *x* that are on the path from *a* to *b. M*2(*a, b*) values can be computed in *O*(*n*^2^) time using the recurrence: *M*2(*a, a*) = *Q*(*a*); for *b* ≠ *a*:

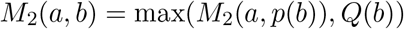

Finally, for each node *a* and a node *b* ≠ *a* where *b* ∈ nodes(*Ta*), we define:

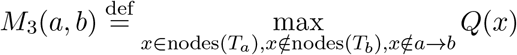

That is, *M*3(*a, b*) is the maximum value of *Q*(*x*) over all nodes *x* that are within *Ta* but not within *Tb* or along the path from *a* to *b*. As an example, in Fig. 11, when calculating *M*3(*r, v*), the node *u* is excluded (because *u* ∈ *Tv*); similarly, the sibling of *r* is also excluded because it is not in *Tr*; on the other hand, *w*1 is included because *w*1 ∈ *Tr, w*1 ∉ *Tv* and *w*1 ∉ *r* → *v*. All *M*3(*a, b*) values can be computed in *O*(*n*^2^) time. This is because there are *O*(*n*^2^) *M*3(*a, b*) values and each value takes *O*(1) time to compute. To see this, we first iterate over each node *b*; then walk the way up to the root from *b*; for each node *a* that is ancestral to *b*, let *a*1 be the child of *a* that is on the path *a* → *b*, and let *a*2 be the sibling of *a*1 but *not* on *a* → *b*. Then *M*3(*a, a*) = −∞. If *a* ≠ *b*,

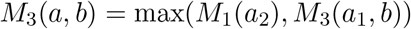

### Constant-time calculation of maximum posterior probability for a single SPR move

We are now ready to show how to calculate the maximum *Q*^′^ value for *T* ^′^ obtained by a single SPR move in *O*(1) time *per SNV site*. We define child(*a, b*) for a node *a* and a node *b* that is a descendant of *a* where child(*a, b*) is the immediate child of *a* that is also ancestral to *b*. For example, in Fig. 11, child(*v*1, *v*2) = *v*. Now consider a single SPR move that prunes a subtree rooted at *u* (*u*’s parent is *v, v*’s parent is *v*1 and *u*’s sibling is *v*2) to form a new node *w* by breaking an edge (*w*1, *w*2) as in Figure 11. With these definitions, we have:

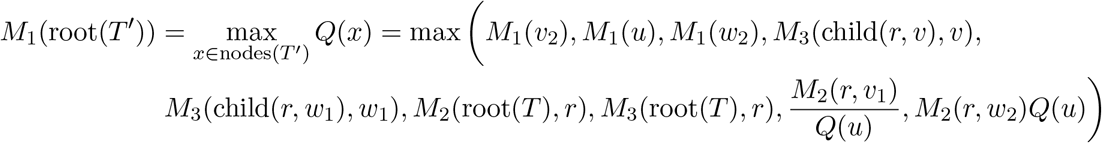

The maximum posterior probability *P* (*T* ^′^) can be computed from *M*1(root(*T* ^′^)) by Equation 2. Here, root(*T* ^′^) is the root node of *T* ^′^. Due to space limitations, we give the detailed algorithm for the SPR local search in the Supplemental Methods (Sect. S3). The correctness of the algorithm is given in the Supplemental Methods (Sect. S4).

### Time analysis

For a fixed SPR operation, the maximum *Q*^′^ values at each site *s* can be computed in *O*(1) time (Equation 3), when the *M*1, *M*2 and *M*3 values are pre-computed for this SPR operation. There are *m* sites. So it takes *O*(*m*) time to compute the maximum posterior probability for a tree in the 1-SPR neighborhood whose size is *O*(*n*^2^). Pre-computing *M*0, *M*2 and *M*3 values takes *O*(*n*^2^) time for each site. Therefore, finding the maximum posterior probability for a single SPR takes *O*(*n*^2^*m*) time in total.

### Branch and bound speedup of SPR local search

Exhaustive search over all the *O*(*n*^2^) trees in the 1-SPR neighborhood can be slow when *n* is large. We now show that a *branch and bound* approach can, in practice, significantly speedup the SPR local search. We use Fig. 11 for illustration. Suppose that we are to prune a subtree that is rooted at *u*. We need to find the best branch (*w*1, *w*2) to regraft for this subtree *Tu*. We let the LCA node of *u* and *w*2 be *r*. Let *w*0 be the child of *r* that is ancestral to *w*2. That is, (*w*1, *w*2) is within 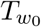. Naïvely, examining all branches under 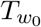 takes *O*(*n*) time because there are *O*(*n*) edges in 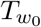. The key idea is that an *upper bound B* of the maximum *Q*^′^ value for all trees that can be obtained by SPR moves that prune *Tu* to *somewhere within* 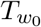 can lead to significant speed up: we can discard *all* SPRs that regraft *Tu* to be within the subtree 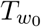 if *B* is no larger than the current maximum *Q*^′^ value we have found so far. Of course, to make the branch and bound approach work, this upper bound needs to be relatively strong and also computable efficiently.

Due to space limitations, we provide the details of the branch and bound in the Supplemental Methods (Sect. S5).

### Genotype calling

Once the optimal tree *Topt* is inferred, genotype calling becomes straightforward. We consider each SNV site *s*, and find the branch *b* of *Topt* to place the single mutation for *s* that maximizes the *Qs*(*v*) value in Equation 2. The cells that are descendants of *b* are the mutants, and the rest are the wild-types. We compute *b* by using the standard trace-back technique in dynamic programming.

## Data Access

ScisTree2 is available on GitHub. ScisTree2 is implemented in C++. We also provide a Python interface. All source code and a tutorial can be accessed at https://github.com/yufengwudcs/ScisTree2. The simulated data and scripts used to reproduce the results presented in this paper are available on Zenodo at https://zenodo.org/records/15620911. The HGSOC dataset can be downloaded from https://doi.org/10.5281/zenodo.5725635.

## Acknowledgments

Research was partly supported by U.S. National Science Foundation grant IIS-1909425 (to Y. Wu). We thank anonymous reviewers and Derek Aguiar for constructive comments on earlier versions of our manuscript.

