## Supplemental Materials for "Large-scale Inference of Cell Lineage Trees and Genotype Calling from Noisy Single-Cell Data Using Efficient Local Search"

**Supplemental Methods** **2**

**Supplemental Fig. S1: Accuracy of trees by neighbor joining with fixed genotypes vs. uncertain genotypes** **16**

**Supplemental Fig. S2: Simulation of adding noise in the data** **17**

**Supplemental Fig. S3: Uncertainty in parameters for HGSOC data analysis** **18**

**Supplemental Fig. S4: Detailed running time of various methods** **19**

**Supplemental Table S1: Speedup by Branch-and-Bound** **20**

Supplemental References . . . . . . . . . . 21

#### Supplemental Methods

#### S1 The probabilistic model of *ScisTree*

*ScisTree2* and *ScisTree* share the same probabilistic model and the overall local search approach. For better understanding of *ScisTree2*, we present the most important aspects of *ScisTree* here: the probabilistic model and the algorithms. For more details, see Wu (2020).

The raw data for *ScisTree* and *ScisTree2* are the sequence reads $D$. Suppose that we are to infer the genotypes $G$. The maximum likelihood estimate $G^a$ is:

$$G^a = \underset{G}{\operatorname{argmax}}\, Pr(D|G)$$

Finding $G^a$ by enumerating $G$ using the above equation directly is infeasible when the number of cells $n$ and the number of sites $m$ are not very small. *CellPhy* computes the likelihood $Pr(D|T)$ by incorporating the uncertainty of $G$ at each leaf of $T$ using a modified version of the well-known Felsenstein's algorithm. One major computational difficulty for *CellPhy*'s likelihood is its complexity: $T$ has branch lengths, and $Pr(D|T)$ needs to (implicitly) sum over all possible alleles at internal nodes. This leads to a relatively complex probabilistic model. This is perhaps the reason why *CellPhy* becomes slow for large number of cells (say $10,000$) despite the fact that *CellPhy* builds on top of RAxML-NG (Stamatakis, 2014), one of the most well-engineered phylogenetic tree inference tools. Moreover, *CellPhy*'s likelihood can not be easily extended to support the IS model.

**The posterior probability model**. *ScisTree* and *ScisTree2* use the *posterior* probability of genotypes $G$ conditional on the given sequence reads $D$ and the IS model:

$$Pr(G|D,I) = \frac{Pr(G,I|D)}{\sum_{G_1} Pr(G_1,I|D)} = \frac{1}{C} Pr(G,I|D) \quad \text{(S1)}$$

where $I$ is the event for the underlying mutations that satisfy the IS model. Note that $C$ is a constant for a fixed $D$. Now,

$$Pr(G, I|D) = Pr(I|G, D)Pr(G|D) = Pr(I|G)Pr(G|D)$$

Here, $Pr(I|G, D) = Pr(I|G)$ because for a fixed $G$, $I$ and $D$ are conditionally independent. If $G$ does not satisfy the IS model, then $Pr(I|G) = 0$. On the other hand, if $G$ satisfies the IS model, it is still possible that the IS model is violated: more than one mutation at a site may not lead to an observable incompatibility with the IS model in $G$. For simplicity, we *assume* that $Pr(I|G) = 1$ if $G$ is compatible with the IS model. Therefore, we can restrict $G$ to only those genotypes that satisfy the IS model. Therefore, Equation S1 is simplified for such $G$:

$$Pr(G|D, I) = \frac{1}{C}Pr(G|D)$$

In the following, for clarity we ignore this constant, and restrict our attention to genotypes that satisfy the IS model. That is, for any $G$ satisfying the IS model, the posterior probability

$$Pr(G|D, I) = Pr(G|D)$$

**Optimization in** ***ScisTree*****.** *ScisTree* and *ScisTree2* aim at finding the genotypes $G^*$ that maximizes the posterior probability where $G^*$ satisfies the infinite sites (IS) model:

$$G^* = \operatorname*{argmax}_{G \in \mathcal{G}^I} Pr(G|D) = \prod_{s=1}^{m} \prod_{c=1}^{n} Pr(G[c, s]|D) \tag{S2}$$

where $c$ refers to a cell, $s$ refers to an SNV site and $\mathcal{G}^I$ is the space of all genotypes that satisfy the IS model. The above equation is due to the assumption of independence among the genotypes of the cells. For a fixed $G$, the probability $Pr(G[c, s]|D)$ in Equation S2 is the *posterior* probability of $G[c, s]$ (the genotype of the cell $c$ at the site $s$ being $G[c, s]$)for $D$) and is computable from the outputs of standard genotype callers.

Recall that we say $G$ satisfy the IS model if there exists a (possibly multifurcating) phylogeny $T$, called *perfect phylogeny* (Gusfield, 1991), where leaves are labeled by the cells (rows) in $G$, and each column (site) $c$ of $G$ labels a single branch of $T$ such that exactly the cells below this branch are the mutants of $c$. Also note that under the IS model, $G^*$ *uniquely* identifies a rooted perfect phylogeny. Thus after obtaining $G^*$, we can use this perfect phylogeny as the underlying cell lineage tree. However, it is known that finding $G^*$ is NP hard (Wu, 2020).

To overcome this computational difficulty, *ScisTree* made the following observation: if the underlying (rooted and binary) tree $T$ is *given*, then we only need to determine the best genotypes conditional on $T$:

$$G^*(T) = \underset{G \in \mathcal{G}^I}{\operatorname{argmax}}\, Pr(G|D,T) \tag{S3}$$

The critical observation is that $G^*(T)$ can be easily determined for any given $T$ in polynomial time using algorithms in Sect. S2. This is because there are only $O(n)$ branches in $T$ for placing the (single) mutation for each site $s$. Once the mutation is placed for a site $s$, the genotypes at $s$ for all cells are determined, and the posterior probabilities of the genotypes can be computed. Thus, for each site, we can simply examine each branch and place the mutation at the branch that gives the largest posterior probability. Since we assume the genotypes of different sites are independent, we can find the optimal genotypes of a site independently of other sites.

Since the underlying tree $T$ is not known, *ScisTree* takes the local search approach to find the optimal binary cell lineage tree $T^*$ and $G^*(T^*)$: (i) construct an initial tree $T_0$, and (ii) iteratively search for a tree $T_m$ by making a single tree rearrangement (say subtree prune and regraft) from the current tree $T_{m-1}$ where $T_m$ gives the largest posterior probability in Eq. S3. Eventually $T_m$ converges to a local optima $T^*$. Then we can determine $G^*(T^*)$ from the optimal $T^*$.

#### S2 Algorithms of *ScisTree*

There are two main aspects of the *ScisTree* algorithm. First, it performs the nearest neighbor interchange (NNI) local search. Here, an NNI operation swaps a subtree rooted at a node $p$ with a subtree rooted at a node $q$ where $q$'s sibling is $p$'s parent. *ScisTree* computes the posterior probability of each tree within the 1-NNI neighborhood. Second, for each tree in the 1-NNI neighborhood, *ScisTree* uses the following algorithm to compute the maximum posterior probability.

Suppose that we are given a rooted binary tree $T$. We want to find the genotypes $G^*$ that satisfy the IS model and maximize $Pr(G^*|D,T)$ where $T$ is the underlying cell lineage tree. We denote $P(G^*,T) = max_{G \in \mathcal{G}^I} Pr(G|D,T)$ as the *maximum* posterior probability of genotypes for $T$. Recall that under the IS model, the set of mutant genotypes at a site $s$ corresponds to a subtree in $T$ and the single mutation occurs on the branch entering the subtree root. We define $P_{s,v}(G,T)$ for a site $s$ and a node $v$ as the posterior probability of all genotypes at site $s$ given that the mutation at $s$ occurs on the branch entering $v$. Since $T$ is fixed, we can enumerate each subtree rooted at node $v$ and compute $P_{s,v}(G,T)$. The maximum probability $P_s(G^*,T)$ at site $s$ is:

$$P_s(G^*,T) = max_{v \in \text{Nodes}(T)} P_{s,v}(G,T) \tag{S4}$$

Here, Nodes($T$) is the set of nodes in $T$. Recall that for each cell $c$ and each site $s$, $M[c,s]$ is equal to the *posterior* probability of the cell $c$ having the *wild-type* genotype (0) at the site $s$. Thus, the posterior probability of allele 0 (respectively 1) at the site $s$ and the cell $c$ is $M[c,s]$ (respectively $1.0 - M[c,s]$). Then, for each node $v$ in $T$:

$$P_{s,v}(G,T) = \prod_{u \in \text{Leaf}(T_v)} (1.0 - M[c(u),s]) \times \prod_{v \notin \text{Leaf}(T_v)} M[c(v),s] \tag{S5}$$

Here $T_v$ refers to the subtree rooted at node $v$. Leaf($T_v$) is the set of leaves of $T_v$. And $c(u)$ is the cell corresponding to a leaf $u$ in $T$.

Computing $P_{s,v}(G,T)$ for each $v$ directly from Equations S4 and S5 would lead to $O(n^2)$ time for each site. A simple observation is that we can apply dynamic programming by taking a bottom-up approach as follows. We define $Q_s(v)$ as the *ratio* of the probability of the genotypes within the subtree $T_v$ being genotype 1 and the probability of these genotypes being genotype 0 at the site $s$. That is (also see the main paper),

$$Q_s(v) = \prod_{c \in \text{taxa}(v)} \frac{1 - M[c,s]}{M[c,s]}$$

Then, we have:

$$P(G^*, T) = \prod_{s=1...m} P_s(G^*, T) = \prod_{s=1...m} \left( [max_{v \in \text{Nodes}(T)} Q_s(v)] \prod_{c=1...n} M[c, s] \right)$$

The algorithm for computing $P_s(G, T)$ based on $Q(v)$ for a single site $s$ for a fixed binary tree $T$ is given below, which has the running time of $O(n)$. Computing $P(G, T)$ involves calculating the product of $P_s(G, T)$ over each of $m$ sites and thus takes $O(mn)$ time.

**Algorithm 1** Maximum probability computation of binary genotypes of a single SNV site $s$

**for** node $v \in T$ in the bottom-up order (i.e., leaves first) **do**
  **if** $v$ is a leaf **then**
    $Q_s(v) \leftarrow \frac{1.0 - M[c(v),s]}{M[c(v),s]}$
  **else**
    Let $v_l$ and $v_r$ being the two children of $v$.
    $Q_s(v) \leftarrow Q_s(v_l) Q_s(v_r)$
  **end if**
**end for**
$P_s(G, T) \leftarrow max_{v \in \text{Nodes}(T)} Q_s(v) * \prod_{c=1...n} M[c, s]$
return $P_s(G, T)$

#### S3 Detailed algorithms for the SPR local search in *ScisTree*

For clarity, we provide detailed algorithms for performing the local SPR search.

First, Algorithm 2 is for *preprocessing*. It computes the values of $M_1, M_2$ and $M_3$ for each site for the original tree $T$ *before* performing any SPR moves. That is, Algorithm 2 only runs once before each iteration of the local search.

**Algorithm 2** Preprocessing step for computing $M_{s,1}, M_{s,2}$ and $M_{s,3}$ values for a specific site $s$ from pre-computed $Q_s(u)$ values.

**for** node $u \in T$ in the bottom-up order (i.e., leaves first) **do**
  **if** $u$ is a leaf **then**
    $M_{s,1}(u) \leftarrow Q_s(u)$ ▷ $M_{s,1}(u)$: the largest $Q_s(v)$ for any $v \in nodes(T_u)$ at $s$.
  **else**
    Let $u_l$ and $u_r$ being the two children of $u$.
    $M_{s,1}(u) \leftarrow \max(M_{s,1}(u_l), M_{s,1}(u_r), Q_s(u))$
  **end if**
**end for**
▷ For a pair of nodes $r$ and $v \in T_r$, $M_{s,2}(r, v)$: the largest $Q_s(u)$ along the path from $r$ to $v$ at $s$.
**for** node $v \in T$ **do**
  $M_{s,2}(v, v) \leftarrow Q_s(v)$
  $r \leftarrow v$
  **while** $r \neq root(T)$ **do**
    $M_{s,2}(p(r), v) \leftarrow \max(M_{s,2}(r, v), Q_s(v))$
    $r \leftarrow p(r)$
  **end while**
**end for**
▷ For a pair of nodes $r$ and $v \in T_r$, $M_{s,3}(r, v)$: the largest $Q_s(u)$ for any node $u$ within $T_r$ but is neither in $T_v$ and nor along the path from $r$ to $v$.
**for** node $v \in T$ **do**
  $M_{s,3}(v, v) \leftarrow -\infty$
  $r \leftarrow v$
  **while** $r \neq root(T)$ **do**
    $w \leftarrow sibling(r)$
    $M_{s,3}(p(r), v) \leftarrow \max(M_{s,3}(r, v), M_{s,1}(w))$
    $r \leftarrow p(r)$
  **end while**
**end for**

Algorithm 3 is to find the maximum posterior probability for a tree $T'$ within 1-SPR neighborhood of the current tree $T$. That is, Algorithm 3 runs for a new tree obtained from some SPR operation.

**Algorithm 3** Finding the tree with the maximum probability within one SPR move from the current tree $T$. Return the maximum posterior probability.

**for** $s = 1 \ldots m$ **do**
  Compute $Q_s(u)$ values for each node $u$ in $T$ and the site $s$ using the probability computation algorithm in the original *ScisTree*.
  Compute $M_{s,1}, M_{s,2}$ and $M_{s,3}$ values for the site $s$ using Algorithm 2.
**end for**
$\triangleright$ Consider each rSPR move involving pruning a subtree $T_u$ and regraft s.t. $T_u$ becomes a sibling of the node $v$
$P_{max} \leftarrow -\infty$
**for** node $u \in T$ **do**
  **for** node $w_2 \in T$ s.t. $w_w \notin T_u$ **do**
    $v_2 \leftarrow sibling(u)$, $v \leftarrow p(u)$, $v_1 \leftarrow p(v)$, $w_1 \leftarrow p(w_2)$
    $P \leftarrow 1$
    **for** $s = 1 \ldots m$ **do**
      $M_A \leftarrow \max(M_{s,1}(v_2), M_{s,1}(u), M_{s,1}(w_2)), \frac{M_{s,2}(r,v_1)}{Q_s(u)}$
      $M_B \leftarrow \max(M_{s,2}(r, w_2)Q_s(u), M_{s,3}(child(r, v), v))$
      $M_C \leftarrow \max(M_{s,3}(child(r, w_1), w_1), M_{s,3}(root(T), r))$
      $P \leftarrow P * max(M_A, M_B, M_C) * \prod_{c=1}^{n} M[c, s]$
    **end for**
    $P_{max} = \max(P_{max}, P)$
  **end for**
**end for**
return $P_{max}$

#### S4 Correctness of Algorithm 3 for evaluating one SPR move for a single site

The key for the correctness of Algorithm 3 is that the $Q$ values at tree nodes before and after the SPR move are highly correlated. This can be seen by carefully analyzing cases for $Q'(x)$ for nodes $x'$ in $T'$ as illustrated in Fig. 11 (main text). $Q'$ refers to the $Q$ values *after* the SPR move. $Q$ and $Q'$ are equal for many nodes, while are different in some other nodes. We have the following cases.

1. $Q'(x) = Q(x)$ for the node $x$ that is *outside* the *cycle* formed by the paths $r \to v$, $r \to w$, $v \to u$ and $w \to u$ corresponding to the SPR move. There are the following cases for $x$.

   (a) $x$ is within a subtree below $v_2, u$ and $w_2$; their maximum $Q$ values are the pre-computed $M_1(v_2), M_1(u)$ and $M_1(w_2)$ respectively (the first three terms in Equation (3) of the main text).

   (b) $x$ is within a subtree whose root $v_r$ is the child of some node on the path $r \to v$ or the path $r \to w$ *but* $v_r$ itself is *not* on these two paths. The maximum $Q$ values for such nodes are $M_3(child(r, v), v)$ and $M_3(child(r, w), w)$ respectively (the 4th and 5th terms in Equation (3) of the main text).

(c) $x$ is along the path from $root(T)$ to $r$, whose maximum $Q$ value is $M_2(root(T), r)$ (the 6th term in Equation (3)).

(d) $x$ is outside the subtree rooted at $r$, whose maximum $Q$ value is the pre-computed $M_3(root(T), r)$ (the 7th term in Equation (3)).

2. $Q'(x) \neq Q(x)$ for a node $x$ where $x$ is *on* the path $r \to v_1$ or the path $r \to w_1$.

(a) $x \in r \to v_1$. $Q'(x)$ is equal to $Q(x)$ *divided* by $Q(u)$ after the SPR (the 8th term in Equation (3)). This is because $Q'$ is for the tree *after* the SPR, where the leaves within $T_u$ are pruned (i.e., *absent* from $T_x$ after the SPR).

(b) $x \in r \to w_1$. $Q'(x)$ is equal to $Q(x)$ *multiplied* by $Q(u)$ (the 9th term in Equation (3)). This is after the SPR, the leaves within $T_u$ are regrafted (i.e., *inserted* into $T_x$ after the SPR).

Algorithm 3 calculates the maximum of these nine cases for each site, which is equal to the maximum $Q'$ values at all nodes in $T'$ after the SPR move. Algorithm 3 then returns the product of these maximum values over all sites. Thus, this algorithm correctly computes the maximum posterior probability $P(T')$ after a specific SPR move.

#### S5 The branch and bound approach

We now describe the details of the branch and bound approach. To simplify the exposition, we focus on one SNV site: the upper bound for multiple sites is the product of the bounds of individual sites. For each node $u$, we consider each *ancestor* $r$ of $u$. We let $w$ be a descendant node of $r$ where $LCA(u, w) = r$. Recall that $LCA(u, w)$ is the lowest common ancestor of $u$ and $w$ in the tree. We consider each $w$ in the *top-down* order. Initially, $w$ is set to $w_0$, the child of $r$ that is *not* ancestral to $u$ (i.e., $w_0$ is the root of the subtree right under $r$ not containing $u$). Then we move downwards from $w_0$ to leaves. Throughout the search, we keep track of the maximal $Q'$ value so far, denoted as $Q'_m$. For each node $w$, we calculate $B(u, w)$, an upper bound on the maximum $Q'$ values among all SPR moves that prune $T_u$ to somewhere *within* $T_w$. Here, $w$ is considered to be within $T_w$. There are two cases in this recursive search.

1. If $B(u, w) \leq Q'_m$, then there are no SPR moves within $T_w$ that gives higher $Q'$ values than

$Q'_m$ and the *entire* subtree $T_w$ can be discarded. That is, any SPR operations that regraft $T_u$ to be inside $T_w$ are ignored. This can lead to significant savings of computation.

2. Otherwise, we calculate the $Q'$ value for the SPR operation that regrafts $T_u$ onto the branch that *enters* $w$ using Algorithm 3. If $Q' > Q'_m$, we let $Q'_m \leftarrow Q'$. Regardless whether $Q'_m$ value is updated, we recursively process the two subtrees of $w$ if $w$ is an internal node, and terminate if $w$ is a leaf.

Obviously, to obtain large speedup, choosing a strong $B(u, w)$ bound is critical. The best $B(u, w)$ bound is the exact maximum posterior probability obtained from trees generated by all SPRs that regraft $T_u$ to be inside $T_w$. But computing this exact maximum will *not* lead to any speed up: calculating this maximum would need to run Algorithm 3 for all possible SPRs within $T_w$; but the branch and bound is meant to avoid such exhaustive calculation in the first place.

In the following, we present an upper bound that is effective in reducing the search space *and* is computable in $O(m)$ time (with appropriate preprocessing). In the following, we present an upper bound on a *single* site that can be computed in constant time. The upper bound of the entire data is the product of the upper bound for all sites.

For two nodes $u, w \in \text{Nodes}(T_r)$ where $LCA(u, w) = r$, we let:

$$\begin{aligned} B(u, w) = \max[&M_1(v_2), M_1(u), M_1(w), M_3(\text{child}(r, v), v), M_3(\text{child}(r, w), w), \\ &M_2(\text{root}(T), r), M_3(\text{root}(T), r), \frac{M_2(r, v_1)}{Q(u)}, M_2(\text{child}(r, w), w)Q(u), M_1(w)Q(u)] \end{aligned} \quad \text{(S6)}$$

Here, $v, v_1, v_2$ and $\text{child}(r, v)$ are the same as in Equation (3) of the main text (also see Fig. 11 in the main text). Different from Equation (3), there is only $w$, but no $w_1$ or $w_2$ in Equation S6. We now argue that $B(u, w)$ is an upper bound on the maximum $Q'$ value for the SPR move that prunes $T_u$ and regraft it to somewhere within $T_w$. Equation S6 is closely related to Equation (3) (the exact maximum $Q'$ for the single SPR move shown in Fig.11 of main text). Intuitively, $B(u, w)$ in Equation S6 is a *relaxed* version of the exact maximal probability of an SPR move in Equation (3): instead of specifying a single branch $(w_1, w_2)$ to regraft, Equation S6 provides an upper bound on the posterior probability for a tree obtained by regrafting to *any* branch within

$T_w$. More specifically, Equation S6 and Equation (3) share the same eight terms. Equation S6 has two terms that are *not* in Equation (3):

1. $M_1(w)$, which is the upper bound on the $Q$ values for nodes inside $T_w$ whose $Q$ values are *not* changed by the SPR.

2. $M_1(w)Q(u)$, which is the upper bound on the $Q$ values for nodes inside $T_w$ whose $Q$ values are *changed* by the SPR.

The other eight terms cover all the $Q$ values outside $T_w$.

For each $u$ and $w$, computing $B(u, w)$ takes constant time. This is because all the $M_1, M_2$ and $M_3$ values have been pre-computed for the current tree. Recall that we may need to compute $B(u, w)$ for each node $w$ in $T$ visited during the top-down recursive search. At each $w$, we may need to run Algorithm 3. Thus, computing $B(u, w)$ takes the same time asymptotically as running Algorithm 3 for a single SPR move. That is, computing the bounds doesn't slow down the tree search asymptotically. In practice, the bounds can significantly reduce the search space. Our experiments show that this branch and bound approach achieves significant speedup for the SPR local search. See Supplemental Table S1 for the empirical results on the performance of the branch and bound approach.

#### S6 Initial tree construction

Similar to *ScisTree*, *ScisTree2* constructs the initial tree using neighbor joining. One key aspect for using neighbor joining is the estimation of the pairwise distance between genotypes of two cells. The original *ScisTree* used the Hamming distance of the called (i.e., fixed) genotypes as the pairwise distance between two cells. *ScisTree2* uses a simple probabilistic pairwise distance which accommodates the uncertainty of genotypes. For two cells $c_1$ and $c_2$, we define the expected Hamming distance $d(c_1, c_2)$ as:

$$d(c_1, c_2) = \sum_{s=1}^{m} [M(c_1, s) * (1 - M(c_2, s)) + (1 - M(c_1, s)) * M(c_2, s)] \quad \text{(S7)}$$

We then use the expected pairwise Hamming distance for running neighbor joining. Our results on simulation data (Supplemental Fig. S1) show that the initial trees constructed from the expected pairwise distance are more accurate than those from fixed genotypes.

#### S7 Calculating the posterior probability of sequence reads simulated by *CellCoal*

The data $D$ simulated by *CellCoal* are in the form of sequence read counts for different alleles at each SNV site for each cell. *CellCoal* calculates the likelihood $Pr(D|G)$. We focus on a single cell and a single SNV site. We now show how to calculate the posterior probability $Pr(G|D)$ from $Pr(D|G)$.

There are four possible alleles, A, T, C and G. We encode these alleles as $0, 1, 2$ and $3$ for the four possible bases (A, T, C and G), where 0 is the wild-type allele. The single-cell genotype $G$ has 10 possible values in *CellCoal*. We use the Bayes formula to calculate the posterior probability $Pr(G|D)$ from the $Pr(D|G)$ values in the VCF files generated by *CellCoal*:

$$Pr(G|D) = \frac{Pr(D|G)Pr(G)}{\sum_{a_1=0}^{3}\sum_{a_2=a_1}^{3} Pr(D|G=\{a_1,a_2\})Pr(G=\{a_1,a_2\})} \tag{S8}$$

The prior genotype probability $Pr(G = \{a_1, a_2\})$ is estimated by the Hardy-Weinberg equilibrium: $Pr(G = \{a_1, a_1\}) = f(a_1)^2$ and $Pr(G = \{a_1, a_2\}) = 2f(a_1)f(a_2)$ $(a_1 \neq a_2)$, where $f(a)$ is the allele frequency of the allele $a$. We assign posterior probability of 0.5 (i.e. the probability of being a wild-type is the same as that of being a mutant) to positions without any reads in the genotype probability matrix.

#### S8 Calculating likelihood and posterior probability from read counts for the HGSOC data

The HGSOC data come with the sequence read counts for the called SNV sites. To calculate the posterior probability of genotypes from these raw read counts, we assume a fixed ADO rate $\theta$ as

0.2 and sequencing error rate $\epsilon$ as 0.01. The likelihood $P(D|G)$ of the reads $D$ at a site for a cell is:

$$\begin{aligned} L(G) &= Pr(D|G=\{a_1,a_2\}) \\ &= (1-\theta)\prod_{i=1}^{N_r} Pr(r_i|G=\{a_1,a_2\}) + \frac{\theta}{2}\prod_{i=1}^{N_r} Pr(r_i|G=\{a_1,-\}) + \frac{\theta}{2}\prod_{i=1}^{N_r} Pr(r_i|G=\{-,a_2\}) \\ &= (1-\theta)\prod_{i=1}^{N_r}\left[\frac{1}{2}Pr(r_i|a_1)+\frac{1}{2}Pr(r_i|a_2)\right] + \frac{\theta}{2}\prod_{i=1}^{N_r} Pr(r_i|a_1) + \frac{\theta}{2}\prod_{i=1}^{N_r} Pr(r_i|a_2) \end{aligned} \quad \text{(S9)}$$

$$Pr(r|a) = \begin{cases} 1-\epsilon, & \text{if } r = a \\ \epsilon, & \text{if } r \neq a \end{cases} \quad \text{(S10)}$$

$$G_{ml} = \underset{G}{\operatorname{argmax}}\, L(G) \quad \text{(S11)}$$

where $G$ is the genotype with two alleles $a_1$ and $a_2$, $r$ is a read and $N_r$ is the number of reads. The likelihood $L(G)$ is to used as the inputs of *CellPhy*. The maximum likelihood genotypes $G_{ml}$ based on the calculated $L(G)$ is used for *HUNTRESS*. The posterior probability of the genotypes for *ScisTree2* can be computed by incorporating a prior on genotype probability using Equation (S8). Allele frequencies are estimated from the maximum likelihood genotypes.

#### S9 Simulation of data by mixing the infinite site model and the finite site model

We want to simulate data where a portion of sites follow the finite sites (FS) model. *CellCoal* offers an option to specify the proportion of sites under the FS model with a relative mutation rate compared to the average mutation rate. However, *CellCoal* does not produce the desired output directly; for instance, it does not explicitly report the number of FS model sites in the simulated data.

To address this limitation, we utilize *CellCoal* to generate data and subsequently merge multiple sites into single sites to emulate the FS model. More specifically, if we aim to generate data with 100 cells, 500 sites, and an FS model proportion of 0.2, we first use *CellCoal* to simulate 10,000 sites under the infinite-site (IS) model, which is significantly more than the desired number of sites. We then retain the first 400 sites unchanged to represent IS model sites. For the remaining 100 FS model sites, we iteratively merge pairs of adjacent IS model sites until the total number of sites is reduced to 500.

#### S10 Robustness of the HGSOC data analysis

There are several aspects that can complicate real data analysis. First, some real data may have more noise than the data we simulated before. Second, there is uncertainty in parameters used by *ScisTree2* and other methods such as priors when analyzing real data. Such parameters may affect the analysis results. In the following, we investigate how robust analysis results from different methods can be on the real HGSOC data.

**Noise in the data**. It is possible that some real data may have higher level of noise than the HGSOC data. To test how robust *ScisTree2* works for data with more noise, we generate semi-simulated data by adding various level of noise into the HGSOC data. We test two ways of adding noises into the HGSOC data.

1. Randomly adding or discarding reads. For each site for a cell, perturb its read count by a random number. The probability of changing the read count is the level of added noise. This is to simulate data with noises in read counts.

2. Randomly masking the genotype probability to be 0.5 (i.e., becomes a missing value) for some site at a cell. Mask rate, the fraction of positions masked to be missing, is the level of added noise. This is to simulate data with large number of missing values.

**Uncertainty in parameters**. To calculate the posterior probability from sequence reads, two parameters are required: the dropout rate and the genotype prior (Equation S9). In real data, there are uncertainty in these parameters. To evaluate the impact of such uncertainties, we conducted tests running *ScisTree2*, *CellPhy*, and *HUNTRESS* on the HGSOC dataset, employing various settings for these two parameters. Specifically, we tested dropout rates of 0.2, 0.5, and 0.75, alongside a non-informative prior that assigns equal probabilities to all genotypes. This resulted in six distinct combinations of dropout rates and priors.

Experiments (Supplemental Fig. S3) show that *ScisTree2* is robust across different parameter values. This indicates that extensive fine-tuning may not be required. We recommend selecting a dropout rate between 0.2 and 0.75.

For genotype priors, we evaluate two approaches: (i) deriving priors from allele frequencies based on Hardy-Weinberg equilibrium and (ii) employing a non-informative prior, where genotype probabilities are set to 0.5. Our results indicate that the option (i) improves the overall inference accuracy. Therefore, for practical applications, we recommend using genotype priors computed from allele frequencies.

### Supplemental Fig. S1: Accuracy of trees by neighbor joining with fixed genotypes vs. uncertain genotypes

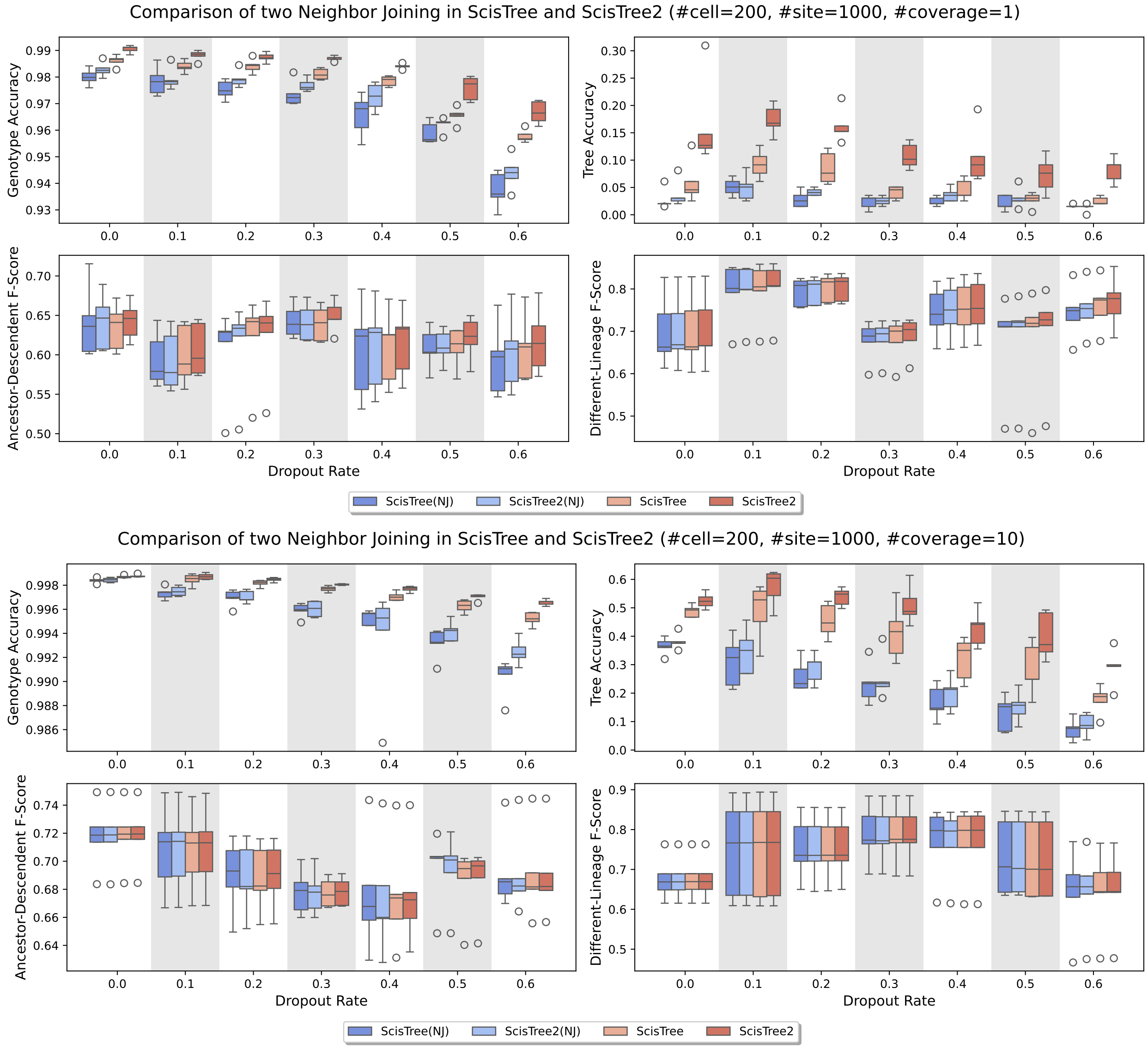

Figure S1: Accuracy comparison of the neighbor joining (NJ) approach with *ScisTree* and *ScisTree2*. Top: low coverage (1x). Bottom: high coverage (10x). X-axis: varying dropout rates. Y-axis: accuracy. NJ performs reasonably well when the data has high coverage and low dropout rate. However, NJ performs worse than *ScisTree2* for data with lower quality.

#### Supplemental Fig. S2: Simulation of adding noise in the data

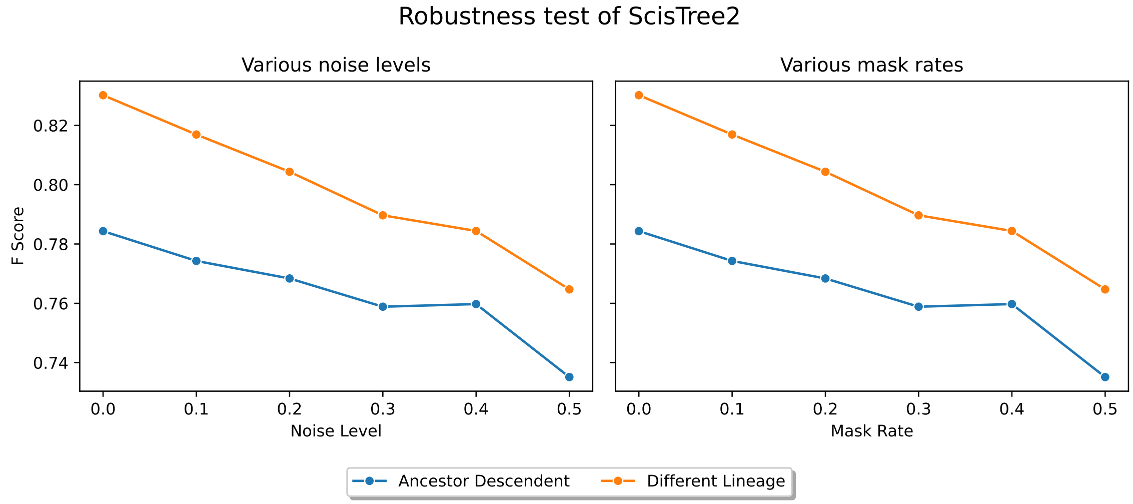

Figure S2: Performance of *ScisTree2* on semi-simulated data by adding noise to the HGSOC data using the approach outlined in the Supplemental Methods. Left: making random changes to the alleles in the read counts for certain percentage of positions. Right: randomly discarding certain fraction of genotype probabilities. Y axis: F score for AD and DL.

#### Supplemental Fig. S3: Uncertainty in parameters for HGSOC data analysis

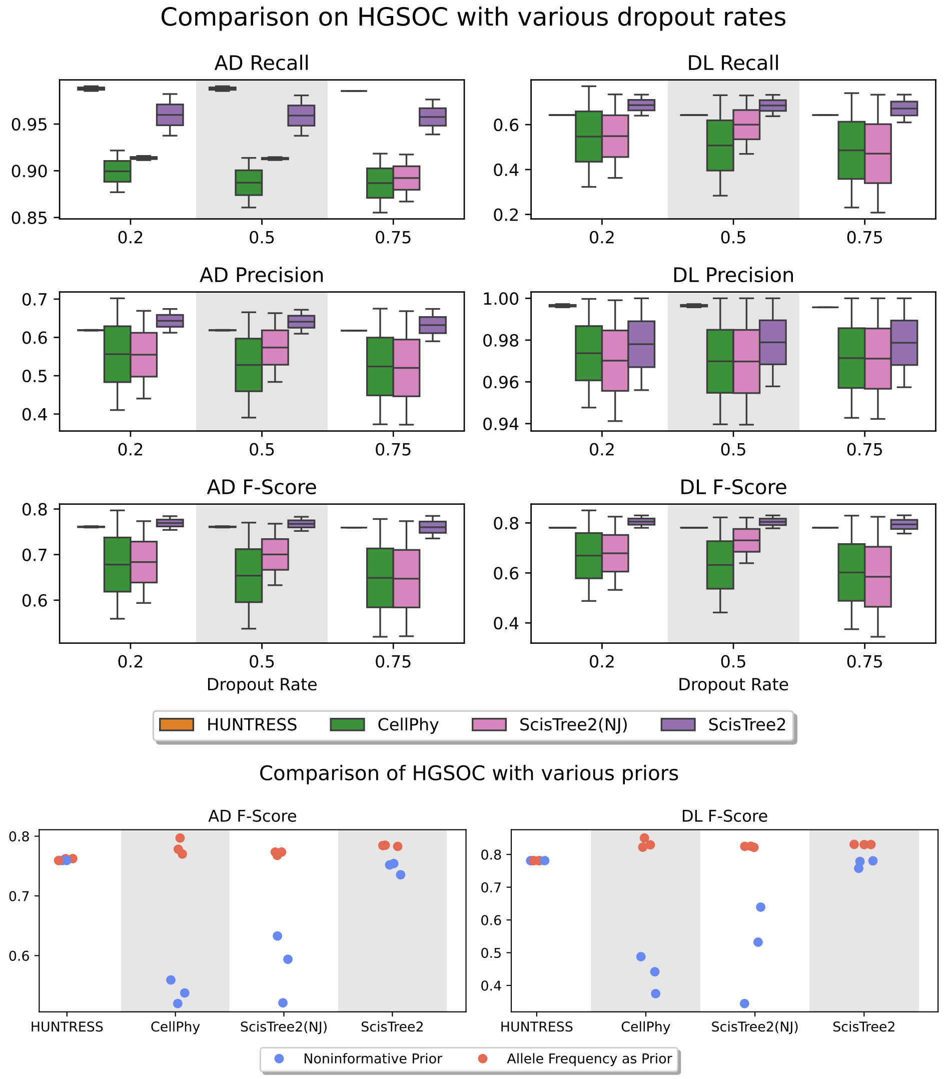

Figure S3: Robustness testing of the effect of parameter uncertainty by varying the ADO rate and genotype priors for the HGSOC data analysis when running *ScisTree2*, *CellPhy* and *HUNTRESS* and NJ (see the Supplemental Methods). AD/DL are used to assess accuracy. Top: varying dropout rates. Bottom: varying the genotype prior: (i) prior calculated from allele frequency by Hardy-Weinberg equilibrium and (ii) non-informative prior. With regards to dropout rates, *ScisTree2* shows robustness across different parameter values.

279 **Supplemental Fig. S4: Detailed running time of various methods**

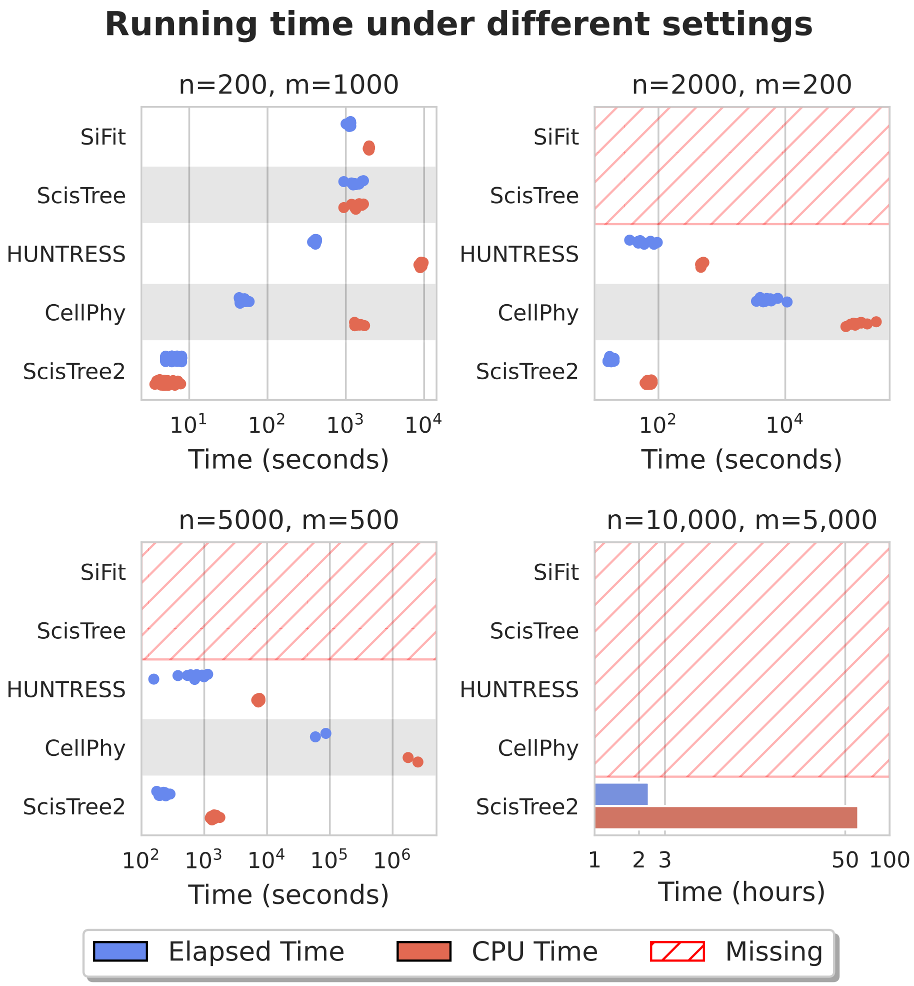

Figure S4: Comparison of the elapsed (user) and CPU running time between *ScisTree2* and other methods on simulated data with the varying numbers of cells and sites. Time is the elapsed time (in hours). 30 threads were used for methods supporting multi-threading. Methods that are too slow are not reported. $n$ is the number of cells. $m$ is the number of sites.

#### Supplemental Table S1: Speedup by Branch-and-Bound

| #Cell | #Site | Elapsed Time(secs) | | | CPU Time(secs) | | |
|---|---|---|---|---|---|---|---|
| | | *ScisTree2 (no BB)* | *ScisTree2* | Speedup | *ScisTree2 (no BB)* | *ScisTree2* | Speedup |
| 100 | 500 | 1.57 | 1.07 | 1.47 | 7.09 | 2.52 | 2.81 |
| 200 | 200 | 3.31 | 1.89 | 1.75 | 11.98 | 3.92 | 3.06 |
| 200 | 400 | 4.14 | 2.35 | 1.76 | 24.13 | 6.67 | 3.62 |
| 200 | 1,000 | 7.43 | 4.13 | 1.80 | 77.07 | 15.32 | 5.03 |
| 200 | 2,000 | 11.71 | 6.46 | 1.81 | 161.15 | 24.48 | 6.58 |
| 500 | 2,500 | 79.41 | 30.77 | 2.58 | 1,575.94 | 140.27 | 11.24 |
| 1,000 | 5,000 | 673.17 | 161.61 | 4.17 | 15,631.39 | 930.22 | 16.80 |

Table S1: A comparison of CPU and elapsed time for *ScisTree2* with and without the branch-and-bound (BB) speedup. All tests were conducted using 30 threads. The branch and bound's effect becomes more evident as the number of cells increases.
